# A *de novo* peroxidase is also a promiscuous yet stereoselective carbene transferase

**DOI:** 10.1101/328484

**Authors:** Richard Stenner, Jack W. Steventon, Annela Seddon, J. L. Ross Anderson

**Affiliations:** School of Biochemistry, University of Bristol, University Walk, Bristol, BS8 1TD, UK; Bristol Centre for Functional Nanomaterials, HH Wills Physics Laboratory, University of Bristol, Tyndall Avenue, Bristol, BS8 1TL, UK; BrisSynBio Synthetic Biology Research Centre, Life Sciences Building, University of Bristol, Tyndall Avenue, Bristol BS8 1TQ, UK; School of Physics, HH Wills Physics Laboratory, University of Bristol, Tyndall Avenue, Bristol, BS8 1TL, UK

## Abstract

By constructing an *in vivo* assembled, catalytically proficient peroxidase, C45, we have recently demonstrated the catalytic potential of simple, *de novo*-designed heme proteins. Here we show that C45’s enzymatic activity extends to the efficient and stereoselective intermolecular transfer of carbenes to olefins, heterocycles, aldehydes and amines. Not only is this the first report of carbene transferase activity in a completely *de novo* protein, but also of enzyme-catalyzed ring expansion of aromatic heterocycles *via* carbene transfer by any enzyme.

## Introduction

Despite the significant advances in protein design, there still remain few examples of *de novo* enzymes constructed from *bona fide*, *de novo* protein scaffolds that both approach the catalytic efficiencies of their natural counterparts and are of potential use in an industrial or biological context^1–9^. This reflects the inherent complexities experienced in the biomolecular design process, where approaches are principally focused on either atomistically-precise redesign of natural proteins to stabilize reaction transition states^1–4^ or imprinting the intrinsic chemical reactivity of cofactors or metal ions on simple, generic protein scaffolds^5–9^; both can be significantly enhanced by implementing powerful, yet randomized directed evolution strategies to hone and optimize incipient function^10,11^. While the latter approach often results in *de novo* proteins that lack a singular structure^6,12,13^, the incorporation of functionally versatile cofactors, such as heme, is proven to facilitate the design and construction of *de novo* proteins and enzymes that recapitulate the function of natural heme-containing proteins in stable, simple and highly mutable tetrahelical chassis (termed maquettes)^14–16^. Since the maquettes are designed from first principles, they lack any natural evolutionary history and the associated functional interdependency that is associated with natural protein scaffolds can be largely circumvented^14^.

We have recently reported the design and construction of a hyperthermostable maquette, C45, that is wholly assembled *in vivo*, hijacking the natural *E. coli* cytochrome *c* maturation system to covalently append heme onto the protein backbone (Fig. 1a)^6^. The covalently-linked heme C of C45 is axially ligated by a histidine side chain at the proximal site and it is likely that a water molecule occupies the distal site, analogous to the ligation state of natural heme-containing peroxidases^17^ and metmyoglobin^18^. Not only does C45 retain the reversible oxygen binding capability of its ancestral maquettes^6,14,15^, but it functions as a promiscuous and catalytically proficient peroxidase, catalyzing the oxidation of small molecules, redox proteins and the oxidative dehalogenation of halogenated phenols with kinetic parameters that match and even surpass those of natural peroxidases^6^.

**Figure 1.**
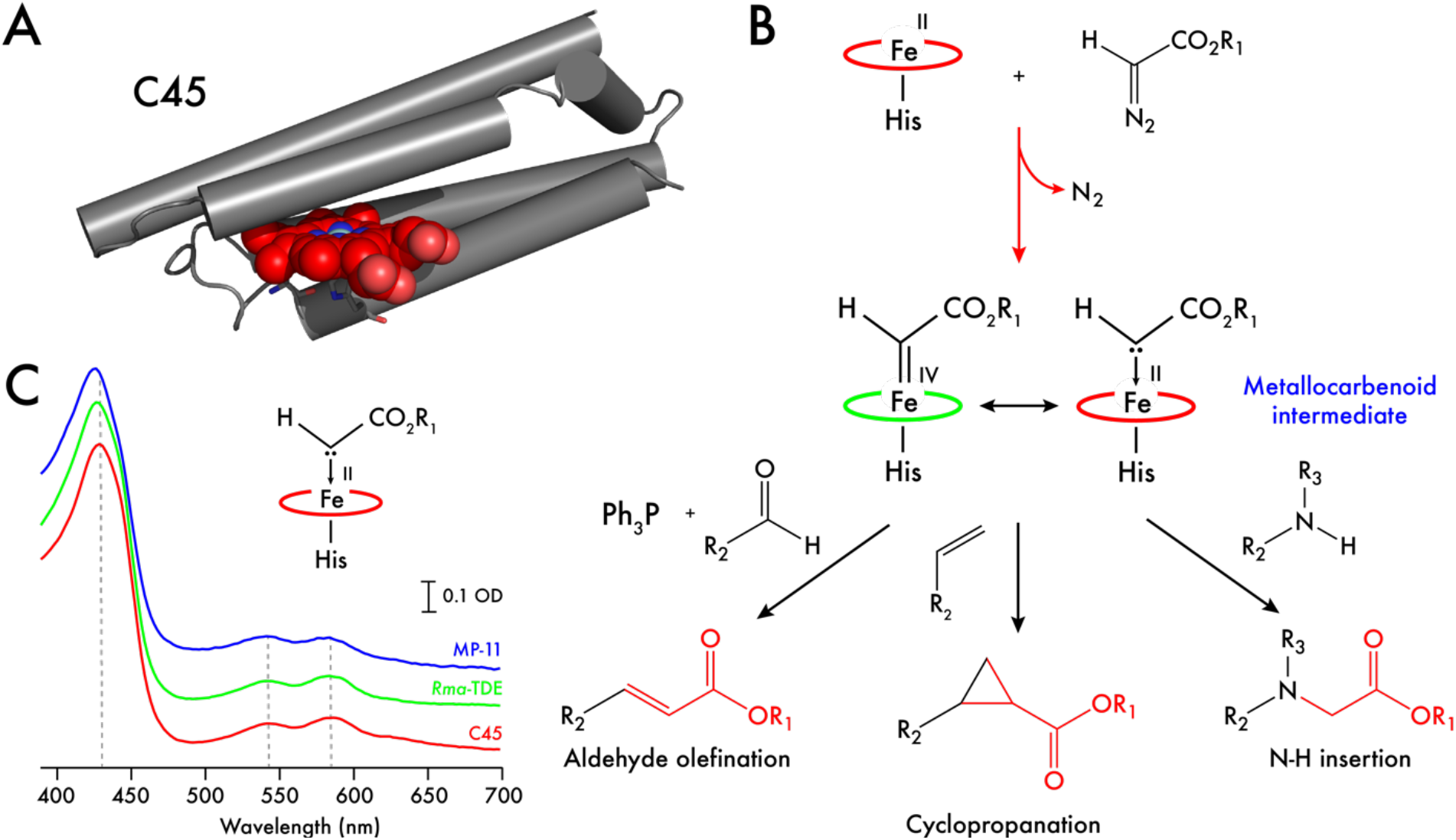
Metallocarbenoid formation and its reactive potential within a *de novo* designed *c*-type cytochrome maquette, C45. **A.** Single snapshot from a 1 μs Molecular Dynamics simulation of the C45 maquette^6^. **B.** Formation and potential reactivity of a heme-based metallocarbenoid intermediate, illustrating aldehyde olefination, olefin cyclopropanation and amine N-H insertion reactions. **C.** UV/visible ethyl diazoacetate-treated C45 (red), engineered *Rma*-TDE (green) and microperoxidase-11 (blue) obtained by rapid mixing experiments in a stopped-flow spectrophotometer. The putative metallocarbenoid species was generated by mixing 2.5 mM ethyl diazoacetate with 7.5 μM ferrous heme protein in 100 mM KCl, 20 mM CHES, pH 8.6 (10% EtOH).

It has been recently demonstrated that several natural heme-containing proteins and enzymes (e.g. cytochromes P450^19^, globins^20^, cytochrome *c*^21^) are capable of accessing many chemistries intrinsic to the heme cofactor, not all of which are essential to life-supporting biological roles. These reported activities are mostly dependent on accessing hypothetical heme-based carbene and nitrene intermediates^22^ analogous to the oxene intermediates, compounds I and II, observed in the catalytic cycles of heme-containing peroxidases and oxygenases^17^. Several groups have now reported several examples of natural and engineered hemoproteins that catalyze cyclopropanations^19,23–30^, C-H insertions^31–34^, carbonyl olefinations^35,36^, N-H insertions^20,37^, C-H amination^38,39^, boron alkylations^40^, aziridinations^41^ and carbon-silicon bond formations^21^, most of which have been exposed to several rounds of directed evolution to improve stereoselectivity and product yield. Excluding the aziridation reactions, an electrophilic metallocarbenoid intermediate (Fig. 1b) is hypothesized to be responsible for carbene transfer to a suitable nucleophile (e.g. olefin)^22,42^ in these reactions. We therefore reasoned that if the metallocarbenoid intermediate could be detected spectroscopically in C45, then the *de novo* enzyme may function as a promiscuous carbene transferase, catalyzing a range of important and challenging organic transformations.

## Results and discussion

To spectroscopically isolate the metallocarbenoid intermediate, we rapidly mixed ferrous C45 with the carbene precursor, ethyl diazoacetate (EDA) at 5 °C in a stopped-flow spectrophotometer. Concomitant with the disappearance of the ferrous C45 spectrum was the appearance of a new spectroscopically distinct species over 60 seconds, with a red-shifted Soret peak at 429 nm and broad Q-bands centered at 543 and 586 nm (Fig. 1c, 2a). Under these conditions, this new spectrum persisted for 1000 seconds with almost no degradation (Fig. 2b), and it is spectroscopically consistent with spectra reported for experimentally-produced carbene:iron porphyrin complexes^43^. For our proposed C45 metallocarbenoid intermediate, the cold conditions on rapid mixing proved essential to spectroscopic observation, as room temperature experiments resulted in spectra indicative of carbene-induced heme degradation, consistent with a mechanism proposed by *Arnold et al*^44^. It was also necessary to employ an ethanol:water mixture to ensure EDA solubility and stability for generating the putative intermediate in the stopped-flow at 5 °C, and we observed an identical, long-lived spectrum at ethanol concentrations between 10 – 50% (SI, Fig. S1). While ethanol concentrations above 50% are generally avoided in buffered protein solutions due to denaturation, small helical bundles such as C45 can readily tolerate such aqueous:organic mixtures^5,6^, retaining structure and catalytic activity. Substituting EDA for benzyl- and *tert*-butyl-diazoacetates (BnDA & ^*t*^BuDA) also resulted in the appearance of intermediates with near identical spectroscopic properties to the EDA-generated C45 species (Fig. 2c), demonstrating the intrinsic flexibility of the active site in accommodating bulky diazoacetate substituents. We were subsequently able to generate and measure mass spectra of these putative intermediates using mild ionization/near native conditions using positive electrospray ionization mass spectrometry (ESI-MS) (SI, Fig. S2). Mass spectra of the EDA-, BnDA- and ^*t*^BuDA-generated metallocarbenoids of C45 exhibited the mass differences expected for the adducts, further confirming the nature of the species generated under these conditions.

**Figure 2.**
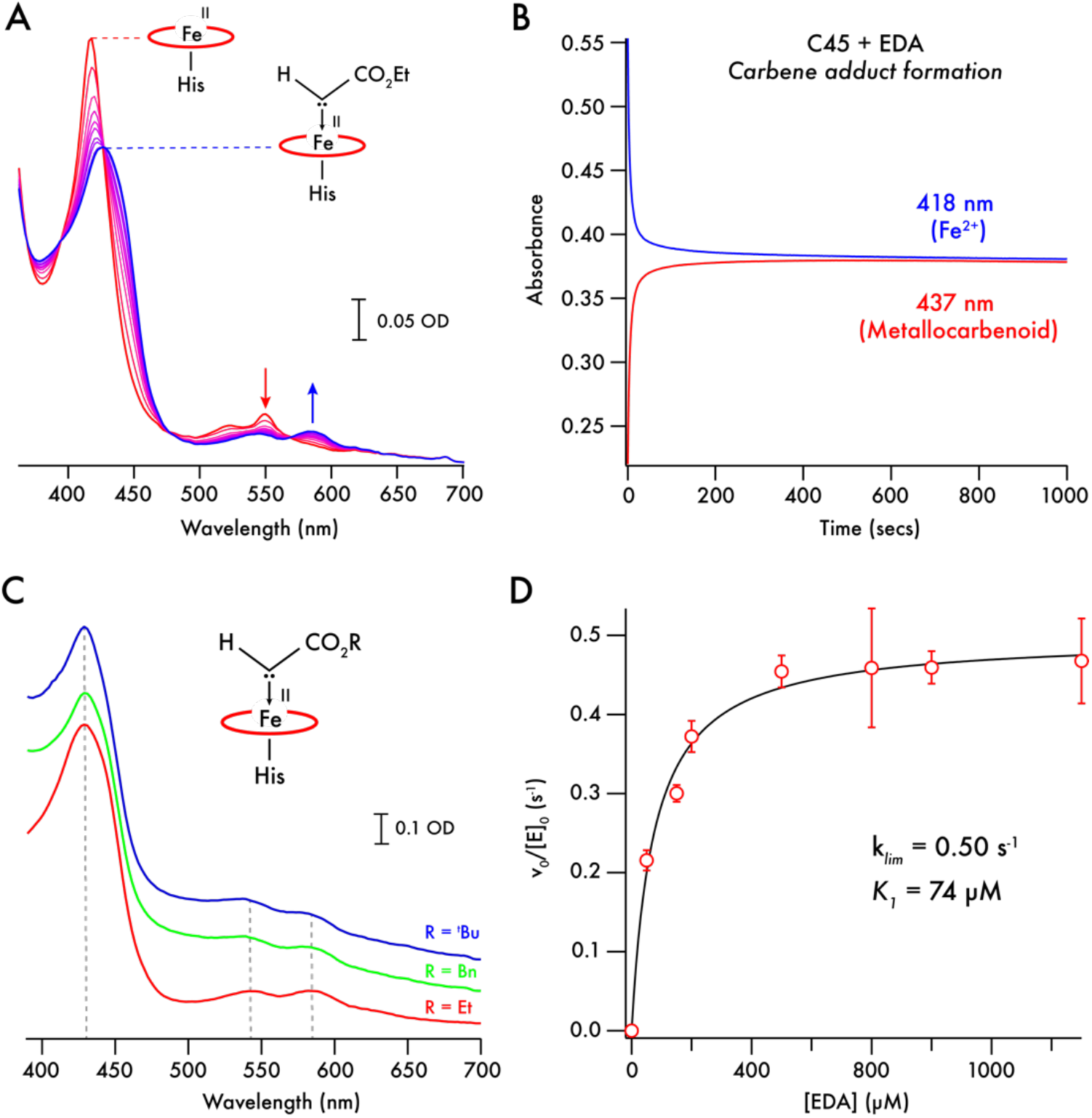
Stability and reactivity of the metallocarbenoid:C45 intermediate. **A.** Time course of electronic spectra recorded following rapid mixing of ferrous C45 (red) with EDA in 10% EtOH at 5 °C. The appearance of the metallocarbenoid intermediate (blue) is concomitant with the disappearance of the ferrous C45 spectrum. Spectra presented were recorded 1, 2, 4, 6, 8, 10, 15, 20, 50 and 100 seconds after mixing. **B.** Metallocarbenoid formation and stability in the absence of styrene substrate. Single wavelength traces represent the time course of ferrous C45 (418 nm, blue, 7.5 μM protein, 10% EtOH) and metallocarbenoid:C45 adduct (437 nm, red) following rapid mixing of ferrous C45 with 500 μM ethyl diazoacetate at 5 °C. Once formed, the metallocarbenoid:C45 adduct persists for the duration of the experiment (1000 seconds). **C.** Electronic spectra of metallocarbenoid intermediates formed between ferrous C45 and ethyl-, benzyl- and *tert*-butyl-diazoacetates following rapid mixing in the stopped-flow apparatus. **D.** EDA-concentration-dependent formation of the C45 metallocarbenoid adduct. Kinetic data were recorded using a stopped-flow spectrophotometer and analyzed as described in the Materials and Methods. The limiting rate constant (*k_lim_*) and *pseudo*-Michaelis constant (*K_1_*) for metallocarbenoid formation are 0.50 s^-1^ and 74 μM respectively.

Under identical conditions to C45, we individually mixed ferrous microperoxidase-11 (MP-11) and an engineered cytochrome *c* from *Rhodothermus marinus*^45^ (*Rma*-TDE) with EDA (Fig. 1c), the latter with an established carbene transferase activity with respect to silanes^45^. Near identical spectra to the putative metallocarbenoid C45 were obtained, though significant differences were apparent between our spectra and those previously reported for the metallocarbenoid complex of *Rma*-TDE^45^. Since our spectra were collected under more rigorously anaerobic conditions than the previous study (stopped-flow spectrophotometer housed in an anaerobic glovebox; < 5 ppm O_2_) and at lower temperature, we postulate that the previously reported UV/visible spectra instead represent an alternative, yet currently unidentified, species. In contrast to C45, *Rma*-TDE exhibits a lag phase during the formation of the putative intermediate, with noticeably slower kinetics compared to C45 under the same experimental conditions. To probe this further, we measured the EDA concentration-dependent kinetics of the putative metallocarbenoid formation for C45 and *Rma*-TDE (Fig. 2d, SI Fig. S3). C45 exhibits both a higher limiting rate constant (*k_lim_*) for metallocarbenoid formation and lower *pseudo*-Michaelis constant (*K_1_*) for EDA (*k_lim_* = 0.50 s^-1^; *K_1_* = 74 μM) compared to *Rma*-TDE (*k_lim_* = 0.14 s^-1^; *K_1_* = 490 μM), demonstrating that not only does C45 bind EDA more rapidly than *Rma*-TDE, but it likely has a significantly higher affinity for the carbene precursor.

We subsequently investigated the ability of C45 to act as an active carbene transferase in the cyclopropanation of styrene, commonly used as an acceptor for heme protein-derived metallocarbenoids^19^. To confirm the identity of the spectroscopically isolated species as the putative metallocarbenoid intermediate, we rapidly mixed ferrous C45 with 100 μM EDA and 3 mM styrene at 5 °C in a stopped-flow spectrophotometer (Fig. 3). Under these conditions, the same putative intermediate spectrum appeared over 60 seconds, but then decayed slowly to the starting ferrous C45 spectrum, consistent with the proposed mechanism of heme-catalyzed carbene transfer to the olefin in which there is no net transfer of electrons from the heme^46–47^. No other spectroscopically distinct species was observed during this experiment. The putative metallocarbenoid species of the engineered *Rma*-TDE displays analogous behavior to C45 in the presence of styrene, with the disappearance of the metallocarbenoid intermediate and concomitant reappearance of the ferrous *Rma*-TDE spectrum (Fig. 3). In contrast, the aerobically-generated species prepared by room temperature mixing of *Rma*-TDE with EDA did not exhibit such behavior, and there was no detectable cyclopropanation activity. Given these data, we assign the new spectroscopically distinct species we describe above to the C45 and *Rma*-TDE metallocarbenoid intermediates. It should be noted that at this time we cannot definitively assign these spectra as either the non-bridging metallocarbenoid species observed by Lewis *et al*^45^ or the porphyrin-bridging species observed by Hayashi *et al*^48^ in the crystal structures of engineered cytochrome *c* or *N*-methylhistidine-ligated myoglobin variant (Mb(H64V,V68A)) respectively. However, given the identical nature of the C45 and *Rma*-TDE spectra, it would seem likely that the spectra obtained represent the non-bridging ligation observed in the *Rma*-TDE Me-EDA crystal structure^45^. Additionally, while the rate of metallocarbenoid intermediate and subsequent product formation appear relatively low in our stopped-flow experiments, it is worth noting that the selected conditions were necessary for maximizing the quantity of intermediate in the stopped-flow apparatus, and that subsequent activity assays were carried out at higher substrate concentrations (both EDA and styrene), higher temperature and lower ethanol concentrations. This would undoubtedly lead to higher reaction rates than those presented in the stopped-flow data here.

**Figure 3.**
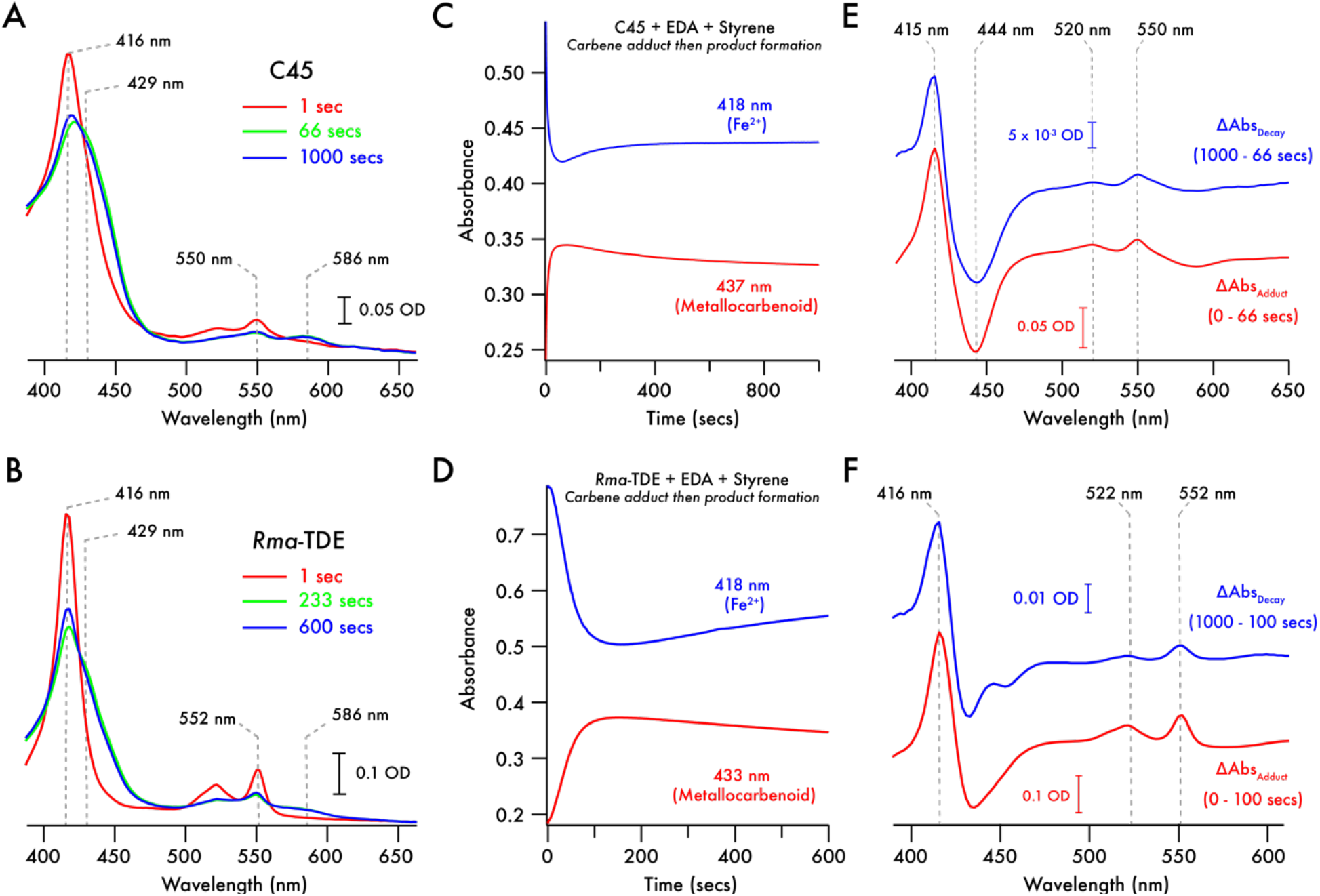
Reactivity of the metallocarbenoid intermediates of C45 and *Rma*-TDE. **A, B.** Electronic spectra of C45 (**A**) and *Rma*-TDE (**B**) recorded after rapid mixing of ferrous heme protein (7.5 μM, red trace) with EDA (500 μM) and styrene (3 mM) at 5 °C in the stopped-flow spectrophotometer. Green traces correspond to spectra recorded at time points where the maximum quantity of metallocarbenoid is accumulated, blue traces correspond to the final spectra recorded in the experiments. **C,D.** Metallocarbenoid formation and decay of C45 (**C**) and *Rma*-TDE (**D**) in the presence of styrene. Single wavelength traces represent the time course of ferrous heme protein (blue traces) (7.5 μM protein) and metallocarbenoid adduct (red) following rapid mixing of ferrous heme protein with 500 μM ethyl diazoacetate and 3 mM styrene at 5 °C. **E,F.** Electronic difference spectra highlighting the spectroscopic changes associated with metallocarbenoid formation and decay for C45 (**E**) and *Rma*-TDE (**F**). The lower, red traces demonstrate the spectroscopic changes that occur during the formation of the metallocarbenoid, which are identical to those observed during its subsequent decay (upper, blue traces). These indicate reformation of the initial ferrous species after carbene insertion and after EDA is exhausted.

Cyclopropanation reactions were initiated at 5 °C under anaerobic conditions and at low ethanol concentrations (5% EtOH) and were subsequently allowed to warm to room temperature. We then analyzed the cyclopropanation activity of C45 and hemin respectively when mixed with EDA and styrene by chiral HPLC and LC-MS. Hemin exhibited little cyclopropanation activity and, under the reaction conditions employed here, only unreacted EDA was observed. However, analysis of the C45-catalyzed reaction revealed the presence of two peaks in the chiral chromatogram at 7.23 and 9.00 minutes, corresponding to the (*R,R*) and (*S,S*) enantiomers of the ethyl 2-phenylcyclopropane-1-carboxylate (Et-CPC) product (SI, Figs. S4 & S5). Following base-hydrolysis of the Et-CPC to the corresponding acid (2-phenylcyclopropane-1-carboxylic acid; CPC) and further LC-MS and chiral HPLC analysis against commercial standards (SI, Figs. S5 & 6), we determined that the C45-catalyzed cyclopropanation of styrene was highly diastereoselective (>99% *de*), occurs at high yield (80.2%, TTN = 802, TOF = 6.68 min^-1^) and exhibits significant enantioselectivity, with an enantiomeric excess (*ee*) of 77% in favor of the (*R,R*) enantiomer. C45 will accept derivatized diazoacetates and *para*-substituted styrenes as substrates for cyclopropanation, producing products with varying stereoselectivities and yields (Fig. 4, SI, Figs. S7-9, Table S1). In particular, C45/EDA catalyzed cyclopropanation of *p*-trifluoromethylstyrene and *p*-methoxystyrene occur with exceptional yield and enantioselectivity for the (*R,R*) product (99.7% yield & 91.4% *ee* for F_3_C-substituted; 90.4% yield & 92.8% *ee* for MeO-substituted). We postulate that the observed stereoselectivity exhibited by C45 results from the intrinsic asymmetry of the protein scaffold, likely through specific positioning of the reactive carbene adduct as proposed by Villarino *et al*^24^, and more recently observed in the crystal structure of the methyl-EDA adduct of the engineered cytochrome *c*^45^. It also offers a means by which we can improve subsequent C45-derived maquettes. In fact, small-scale screening of a C45 library generated by successive generations of error-prone PCR and originally created for improved peroxidase activity revealed a variant of C45 (AP3.2) with switched and remarkably high enantioselectivity towards the (*S,S*) product (*ee* = 99%) (SI, Fig. S10).

**Figure 4.**
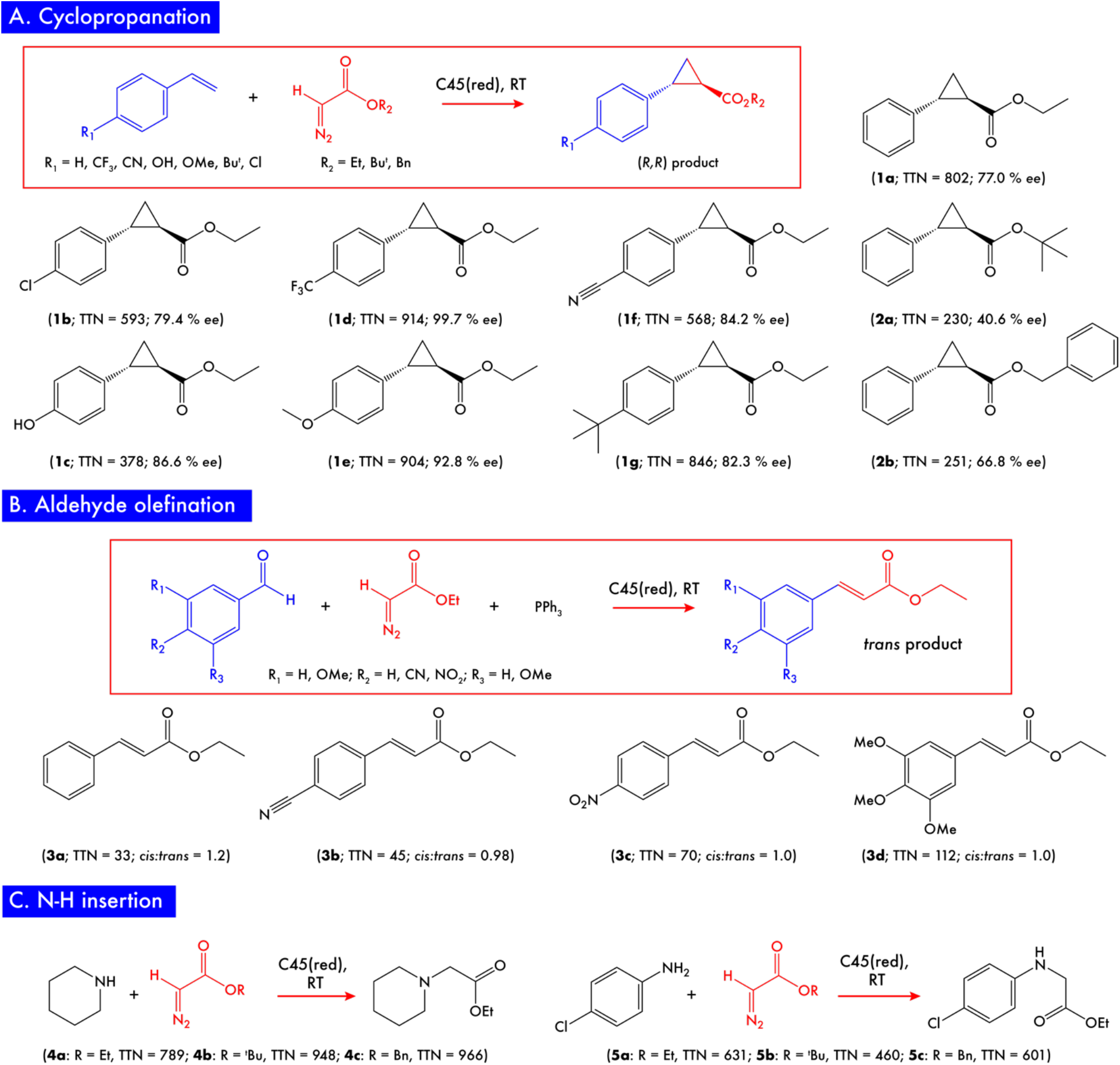
Carbene transferase activity of C45. **A.** Cyclopropanation of substituted styrenes catalyzed by C45. Total turnover numbers (TTN) and enantiomeric excesses (% *ee*) for each combination of ferrous C45 with para-substituted styrenes and functionalized diazoacetates. Only the (*R,R*) cyclopropanated product is displayed in the reaction scheme, representing the dominant product in all cases. All reactions were carried out with 0.1% catalyst loading (10 μM C45) at the following concentrations of reagents: 10 mM sodium dithionite, 10 mM diazo compound, and 30 mM substituted styrene in 100 mM KCl, 20 mM CHES, pH 8.6, 5% EtOH. **B.** Olefination of substituted benzaldehydes catalyzed by C45. Only the *trans* product is displayed in the reaction scheme, though almost equal quantities of the *cis* product are also produced in the C45-catalyzed reactions. All reactions were carried out with 0.1% catalyst loading (10 μM C45) at the following concentrations of reagents: 10 mM sodium dithionite, 10 mM PPh_3_, 10 mM ethyl diazoacetate, and 30 mM substituted benzaldehyde in 100 mM KCl, 20 mM CHES, pH 8.6, 5% EtOH. **C.** N-H insertion of primary and secondary amines catalyzed by C45. Only the monofunctionalized product of the *p*-chloroaniline insertion reaction shown, and the corresponding TTN is calculated based on the yield of both mono- and di-substituted products. All reactions were carried out with 0.1% catalyst loading (10 μM C45) at the following concentrations of reagents: 10 mM sodium dithionite, 10 mM ethyl diazoacetate, and 30 mM amine substrate in 100 mM KCl, 20 mM CHES, pH 8.6, 5% EtOH.

To probe the flipped stereoselectivity and reduced yield exhibited by AP3.2, we purified and characterized the protein both spectroscopically and functionally. AP3.2 exhibits near identical UV/visible ferric and ferrous spectra to C45^6^ (SI, Fig. S11a), and a CD spectrum consistent with that expected for a 4-helix bundle maquette^6,14,16^ (SI, Fig. S11b), albeit with reduced secondary structure compared to C45. The eight mutations to C45 (F11Y/G39S/D48Y/F53S/F83S/G109A/F132S/E133G) destabilize the protein scaffold, with AP3.2 exhibiting a 37 °C drop in the thermal melting transition (*T_m_* = 49 °C) in comparison to C45^6^ (SI, Fig. S11c). Given that three of these substitutions replace core, hydrophobic phenylalanines in helical positions with the hydrophilic serine, this is not surprising, especially given that serine has a significantly lower helical propensity than alanine^49^. Mapping the AP3.2 mutations onto our computationally generated C45 model^6^ (SI, Fig. S11d) does not immediately reveal an obvious explanation for the pronounced change in stereoselectivity we observe, as only the F53S mutation is positioned near to the carbene binding site at the distal face of the heme. To examine reactivity in more detail, we generated the AP3.2 metallocarbenoid intermediate by rapid mixing under the same conditions as described for C45, and we observed an identical intermediate spectrum that accumulates on a comparable timescale to C45 (SI, Fig. S12) and, in the presence of styrene, decays at a comparable rate to C45 with identical associated spectroscopic changes (SI Fig. S13a-c). To test the possibility of an electronic effect playing a role in the suppressed activity of AP3.2, we used redox potentiometry to measure the AP3.2 heme redox potential (*E_m_*) (SI, Fig. S13d). AP3.2 exhibits a 26 mV negative shift in redox potential with respect to C45 (C45 *E_m_* = −176 mV^6^; AP3.2 *E_m_* = −202 mV), indicating slight stabilization of the ferric form, possibly due to increased solvent accessibility at the heme^50^. While this shift also indicates that the extent of electron donation to the bound carbene is indeed altered following the mutations, it is relatively modest in comparison to those observed after heme axial ligand substitution in engineered P450s^50–51^. In these cases, the correlation between large redox potential changes and the specific reactivity of the metallocarbenoid is currently unclear, at least experimentally, and increased reactivity resulting from a more electrophilic carbene adduct would be expected to correlate with a more electron withdrawing, higher redox potential heme^42^. Furthermore, we identified another C45 variant (APR1) from the library with similarly altered redox potential (APR1 *E_m_* = −197 mV; Δ*E_m_* = −21 mV), but with negligible difference in yield or enantioselectivity (yield = 85.6%, TTN = 856, *ee*(*R,R*) = 78.0%; SI Fig. S14), therefore highlighting that such small differences in *E_m_* likely contribute little to the overall yield of product under these reaction conditions. As is the case for many enzymes improved by random laboratory evolution methodologies^51^, it is therefore not unambiguously clear why this altered reactivity and enantioselectivity occurs, and a currently uncharacterized combination of static and dynamic processes likely play a role in the observed catalytic and stereoselective changes. It does, however, demonstrate well the mutability and catalytic potential of the maquette scaffold and will be explored in more depth in future work.

Since there are now several examples of engineered and natural proteins capable of catalyzing carbene transfer, we wished to benchmark the activity of C45 against some notable and well-characterized carbene transferases. To this end, we directly compared the C45-catalysed cyclopropanation of styrene with EDA against the corresponding reactions with *Rma*-TDE^21,45^ and a double mutant of sperm whale myoglobin, Mb(H64V,V68A)^25^ (SI Figs. S15 & S16, Table S2). While *Rma*-TDE was engineered towards silane alkylations^21^, it forms a structurally characterized carbene adduct and is a *c*-type cytochrome, rendering it the most spectroscopically similar carbene transferase to C45 in the literature. In contrast, the cyclopropanation activity of Mb(H64V,V68A), and other Mb variants, towards styrene has been extensively studied and is both high yielding and highly enantioselective for the (*S,S*) product^25^. Under identical conditions to the reactions described above for C45, we confirmed that both proteins catalyze the cyclopropanation of styrene, with Mb(H64V,V68A) exhibiting >99% *ee* for the (*S,S*) product (yield = 95.9%, TTN = 959, TOF = 7.99 min^-1^) as *per* the literature^25^, and *Rma*-TDE exhibiting 70.6%*ee* for the (*R,R*) product (yield = 73.5%, TTN = 735, TOF = 6.12 min^-1^). While the yield from the C45-catalysed reaction (80.2%) does not surpass that of Mb(H64V,V68A), it is among the highest yields observed for the catalytic cyclopropanation of styrene by EDA, surpassing those exhibited by almost all reported P450^19,25–29^ and LmrR^24^ variants, as well as several engineered myoglobins^23,25,29^. In addition, *Rma*-TDE exhibits a relatively high yield and enantioselectivity for styrene cyclopropanation despite being engineered for carbene insertion into Si-H bonds.

With stereoselective carbene transferase activity firmly established for the cyclopropanation of styrene by C45, we explored the substrate promiscuity of C45 towards similar reactions resulting in N-H insertions and carbonyl olefinations. Following reaction conditions identical to those described above for cyclopropanation and using HPLC and LC-MS to identify insertion products, we observed the C45-catalyzed insertion of EDA/^*t*^BuDA/BnDA-derived carbene into the N-H bonds of *para*-chloroaniline and piperidine (Fig. 4, SI, Figs. S17-21, Table S3), with single and double insertion products resulting from carbene transfer to the primary amine of *para*-chloroaniline. Notably, C45 achieves very high yields for the insertion of ^*t*^BuDA- and BnDA-derived carbenes into the piperidine N-H bond, with 94.8% and 96.6% respectively (TTN = 948 for ^*t*^BuDA, TTN = 966 for BnDA). For carbonyl olefination, the addition of triphenylphosphine to the reaction mixture is necessary to facilitate the formation of an ylide that, through the generation of an oxaphosphetane intermediate, spontaneously collapses to the product^35,36^. We investigated the olefination of four potential aldehyde substrates: benzaldehyde, *p*-nitrobenzaldehyde, *p*-cyanobenzaldehyde and 3,4,5-trimethoxybenzaldehyde. After overnight reactions were performed, the products were again analyzed by HPLC and LC-MS, and though yields and TTNs were typically low, they were consistent with values reported for several myoglobin variants (Fig. 4, SI, Figs. S22-24, Table S4)^36^. Interestingly, C45 did not exhibit discernible diastereoselectivity for carbonyl olefination, possibly indicating that the ylide is released from the protein prior to rearrangement and final product formation. Nevertheless, the ability to catalyze these reactions further illustrates the utility of C45 as a general carbene transferase enzyme.

Ring expansion reactions are exceptionally useful in organic synthesis as they provide a reliable and facile method for acquiring large, expanded ring systems^52,53^. Though currently underused, the homologous ring expansion of nitrogen-containing heteroaromatics could be of considerable use in the synthesis of pharmaceuticals and natural products. While both the Buchner ring expansion^54^ and Ciamician-Dennstedt reaction^55^ proceed *via* a cyclopropane-containing bicyclic system that is subsequently ring-opened, the ring opening during the latter reaction is facilitated by the expulsion of a halogen leaving group. We therefore hypothesized that with a halogen-substituted carbene precursor, C45 would catalyze the cyclopropanation of a nitrogen-containing heteroaromatic to produce a cyclopropane-containing bicyclic system susceptible to spontaneous rearrangement (Fig. 5a). Using the same reaction conditions as those described for C45-catalyzed styrene cyclopropanation, we investigated whether C45 could catalyze the ring expansion of pyrrole with ethyl-2-bromo-2-diazoacetate^56^ as the carbene precursor. After 2 hours, the reaction was analyzed using HPLC and LC-MS, confirming the production of the ring-expanded product, ethyl nicotinate at 69.4% yield (Fig. 5b, SI Figs. S25 & S26). Subsequent base-catalyzed hydrolysis of the product yields the nicotinamide and NAD(P)H precursor niacin, raising the possibility of engineering a life-sustaining, artificial biochemical pathway from pyrrole to nicotinamide reliant on the *in vivo* activity of C45 or a related *de novo* designed enzyme. To the best of our knowledge, this is the first example of any enzyme - natural, engineered or *de novo* - that is capable of catalyzing homologous ring expansion reactions *via* a carbene transfer mechanism.

**Figure 5.**
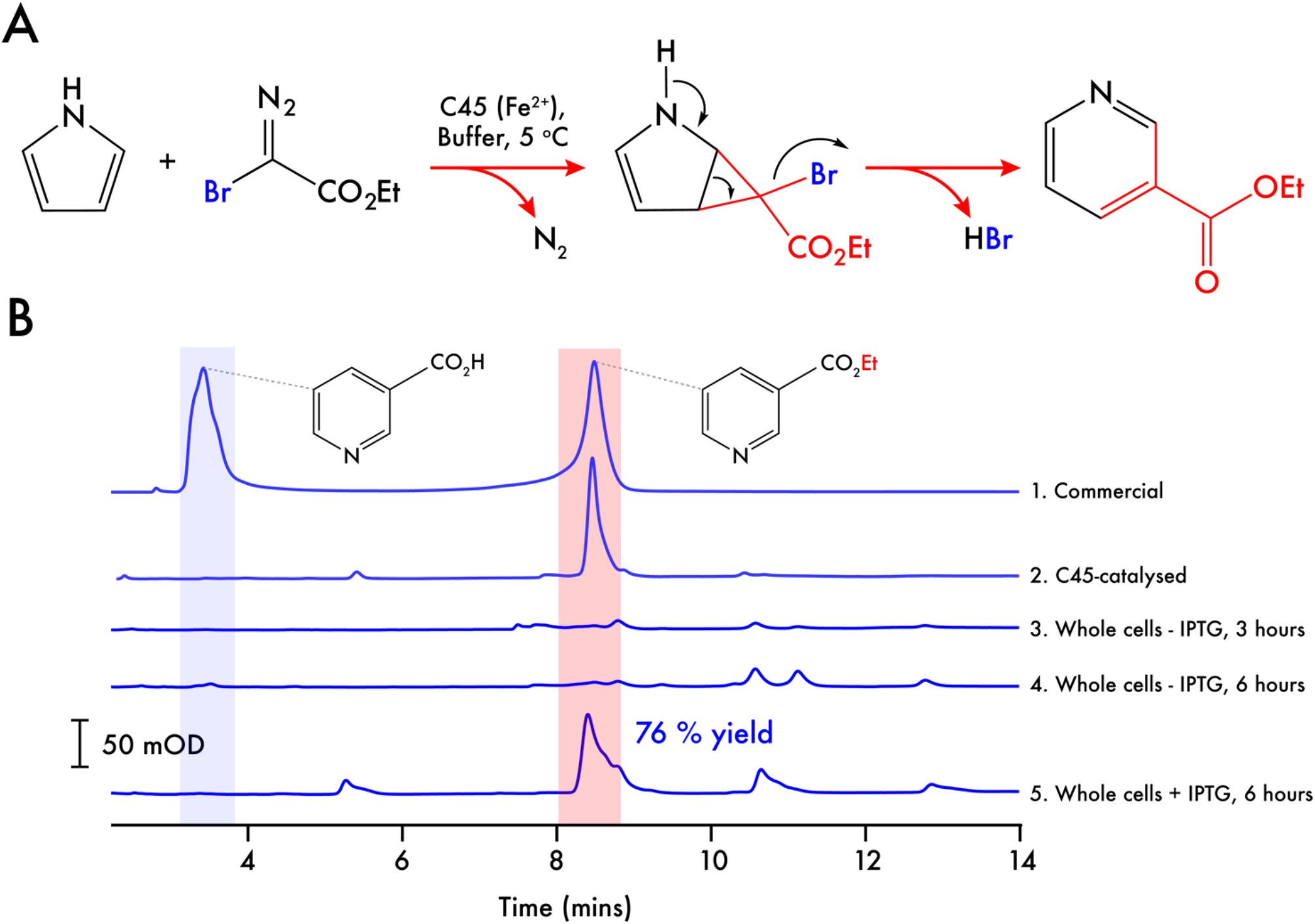
Heteroaromatic ring expansion catalyzed by C45. **A.** Reaction scheme for the ring expansion strategy using ethyl 2-bromo-2-diazoacetate, pyrrole and ferrous C45. Following carbene transfer to the pyrrole, spontaneous rearrangement of the bicyclic ring system leads to elimination of HBr and formation of a 6-membered pyridine ring. **B.** C18 reversed phase HPLC traces of the C45-catalyzed ring expansion of pyrrole to ethyl nicotinate. Traces 1 & 2 show the C45-catalysed ring expansion compared to a partially hydrolyzed commercial standard of ethyl nicotinate. The ring expansion was carried out with 1% catalyst loading (10 μM C45) at the following concentrations of reagents: 10 mM sodium dithionite, 1 mM ethyl 2-bromo-2-diazoacetate, and 10 mM pyrrole in 100 mM KCl, 20 mM CHES, pH 8.6, 5% EtOH. Traces 3,4 & 5 show the results of incubating whole cells containing the C45 expression vector and pEC86 harboring the *E. coli* cytochrome *c* maturation apparatus. Traces 3 & 4 represent reactions between whole cells, pyrrole and ethyl 2-bromo-2-diazoacetate at 3 and 6 hours after inoculation and in the absence of the inducer, IPTG. Trace 5 represents the reaction with C45-expressing whole cells, pyrrole and ethyl 2-bromo-2-diazoacetate. In this case, the cells were grown for 3 hours, induced with 1 mM IPTG and C45 was expressed for a further 3 hours prior to use in the whole cell transformation. Reaction conditions are fully described in the Materials and Methods section.

To further explore the possibility of employing C45 in a new, essential and life-sustaining pathway from pyrrole to the pyridine nucleotides, we examined the ring-expansion of pyrrole to ethyl nicotinate in living *E. coli* cells. Using an established procedure for carbene transferase activity under such conditions^26–28^, we tested the ability of *E. coli* bearing the C45-expression plasmid and pEC86 (harboring the *c*-type cytochrome maturation apparatus)^57^ to perform the ring expansion reaction of pyrrole with ethyl-2-bromo-2-diazoacetate under anaerobic conditions (Fig. 5b). Since the maturation apparatus is constitutively expressed and results in the production of several heme-containing membrane proteins (CcmC, CcmE & CcmF)^58,59^, it was vital to establish whether any intrinsic ring expansion activity was detectable in the presence of these proteins. Whole cells that had been grown for 3 and 6 hours in the absence of the inducer, IPTG, had barely detectable ring-expansion activity (Fig. 5b). In contrast, cells that were grown for 3 hours, induced with IPTG and grown for a further 3 hours displayed significant ethyl nicotinate formation with a total yield of 76% (Fig. 5b), indicating that under these conditions and *in vivo*, C45 exhibits the catalytic ring-expansion activity. It also indicates that both pyrrole and ethyl-2-bromo-2-diazoacetate are able to cross the outer membrane and access the periplasmically-located C45. Interestingly, despite the production of ethyl nicotinate by the cells, we did not observe an increase in the quantity of niacin in the extract, indicating that *E. coli* possibly lacks - or does not produce an appreciable quantity of - an endogenous periplasmic esterase capable of efficiently hydrolyzing the product. We therefore tested a recombinant *Bacillus subtilis* esterase^60^ expressed in, and purified from, *E. coli* for ethyl nicotinate hydrolysis activity. Incubation of this esterase with ethyl nicotinate under near-physiological conditions results in the production of niacin (SI, Fig. S27), thus highlighting another functional part in a potential novel biosynthetic pathway from pyrrole to the pyridine nucleotides through niacin.

To this end, we speculate that it is possible to make C45 and the *Bacillus* esterase essential to NAD+ biosynthesis. *E. coli* synthesizes NAD+ through two pathways that both involve the production of NaMN (SI, Fig. S28), then NaAD prior to NAD+ formation^61–64^: the *de novo* pathway, using *L*-aspartate as a starting material to produce quinolate which is a NaMN precursor^62^; the pyridine ring salvage pathway where exogenous niacin or nicotinamide are utilized instead as NaMN precursors, representing the favored pathway when niacin and nicotinamide are abundant in the environment^64^. Knocking out a key enzyme in the aspartate pathway - e.g. nicotinate-nucleotide diphosphorylase (NadC) - will provide an auxotrophic strain of *E. coli* that, when expressing both C45 and a suitable periplasmically-directed esterase (using a signal sequence such as malE as for C45^6^) in nicotinamide and niacin-lacking media, should be capable of converting pyrrole and ethyl-2-bromo-2-diazoacetate to niacin and ultimately NAD+, recovering the deleterious phenotype. Currently, production of such an *E. coli* strain is outwith the scope of this study, but this strategy highlights the mechanism by which we can tractably and rationally engineer the metabolism of a bacterium to rely on a *de novo* enzyme to sustain an essential pathway.

## Conclusions

With this work, we have showcased the exceptionally diverse functionality available in this simple, *de novo* designed heme protein, and demonstrated that the carbene transferase activity intrinsic to the scaffold compares very favorably to that reported for most engineered natural proteins. The formation of the metallocarbenoid intermediate at the C45 heme facilitates not only the high-yielding cyclopropanation of styrene and its derivatives, but also extends to the insertion of carbenes into N-H bonds, the olefination of carbenes and the first description of enzyme-catalyzed ring expansion of a nitrogen heterocycle. This demonstrates that abiological function is intrinsic to these *de novo* designed maquettes, and that the higher complexity of natural protein frameworks, such as the globins and cytochromes P450, is not necessary to support reactivity of this type. Given the altered product profile exhibited by AP3.2, we have also demonstrated the flexibility of the maquette framework towards substantial mutational change, profoundly affecting the stereoselectivity of the *de novo* enzyme. This indicates that further yield enhancements and alternative stereoselectivities for these types of reaction are undoubtedly achievable through directed evolution, consistent with the studies reported for the maquette’s natural counterparts^19,21,31^. Furthermore, there are many beneficial features of C45 that compare favorably against the work described by others with natural enzymes: C45 is expressed at high levels in *E. coli*, and is functionally assembled *in vivo*^6^, eliminating the requirement for the *in vitro* incorporation of abiological metalloporphyrins^31^ and facilitating *in vivo* function^26^; C45 is thermally stable and displays excellent tolerance in organic solvents^6^, thereby facilitating its use as a homogenous catalyst in aqueous:organic mixtures. In conclusion, this work demonstrates the biocatalytic potential and reactive promiscuity that maquettes possess, and future work will be concerned with improving the substrate scope and catalytic performance of these novel, *de novo* enzymes. It also reinforces our previous work^6^, demonstrating that maquettes and related *de novo* proteins are much more than just lab curiosities or ornaments, and are instead powerful, green catalysts that can play a valuable role in facilitating challenging organic transformations.

## Experimental details

### General & molecular biology, protein expression and purification

All chemicals were purchased from either Sigma or Fisher Scientific and cloned synthetic genes for *Rma*-TDE and Mb(H64V/V68A) were purchased from Eurofins Genomics. Protein expression and purification in *E. coli* T7 Express (NEB) from the pSHT (for periplasmic expression) or pET45b(+) (for cytoplasmic expression of Mb(H64V/V68A)) vectors were carried out as described previously^6^. Error prone PCR (epPCR) libraries of C45, with a mutation rate of 2-3 amino acid mutations, were produced using the GeneMorph II EZClone Domain Mutagenesis Kit. The mutation rate was determined by randomly selecting and sequencing 10 colonies after transforming the PCR products into *E. coli* T7 Express cells. This mutation rate was achieved with 18.7 ng of target DNA (C45 coding sequence). AP3.2 and APR1 were randomly selected variants from the epPCR C45 library. Each variant was expressed and purified as previously described for C45^6^, and the purified proteins were used in the cyclopropanation assays.

### Synthesis of ethyl 2-bromo-2-diazoacetate

Ethyl 2-bromo-2-diazoacetate was synthesized as previously reported^56^. Briefly, *N*-bromosuccinimide (26 mmol) was added to a solution of EDA (20 mmol) and 1,8-Diazabicyclo(5.4.0)undec-7-ene (28 mmol) in CH_2_Cl_2_ (5.0 mL) at 0 °C, and the reaction mixture was stirred for 10 minutes. The crude reaction mixture was washed with cold Na_2_S_2_O_3_ (aqueous, 3 x 5 mL) and quickly filtered through a silica column (cold CH_2_Cl_2_). Cold CH_2_Cl_2_ was added to bring the final volume up to 2.5 mL (~40 mM).

### Stopped-flow spectrophotometry

Stopped-flow kinetics were conducted using a SX20 stopped-flow spectrophotometer (Applied Photophysics) housed in an anaerobic glovebox under N_2_ ([O_2_] < 5 ppm; Belle Technology). In the initial experiments, a solution containing a known concentration of reduced C45 or AP3.2 (15 μM, 100 mM KCl, 20 mM CHES, pH 8.6, reduced with Na_2_S_2_O_4_) was placed in one syringe and a 5 mM solution of a carbene precursor (either EDA, ^*t*^BuDA or BnDA) in ethanol (20-100%) was placed in the second syringe. 50 μL from each syringe was simultaneously injected into a mixing chamber and the progression of the reaction was monitored spectroscopically at 5 °C and 25 °C, over the course of 180-1000 seconds, to examine the metallocarbenoid formation and porphyrin degradation pathway respectively. The formation of the metallocarbenoid was monitored by following the time-course profiles at 417 nm and 428-437 nm, the Soret peak for reduced C45/AP3.2/*Rma*-TDE and the metallocarbenoid intermediate respectively. Final concentrations were 7.5 μM ferrous C45/AP3.2/*Rma*-TDE and 2.5 mM EDA/^*t*^BuDA/BnDA (10-50% ethanol). The kinetics of the formation of the metallocarbenoid for C45/*Rma*-TDE were determined using the same conditions outlined above but using varying concentrations of EDA (50 μM – 1.5 mM). The rate constants were calculated at varying substrate concentrations, plotted and then the data were fitted using the following equation for reversible formation of an unobserved tetrahedral intermediate followed by irreversible removal of dinitrogen to the stable metallocarbenoid species:

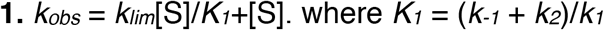

Additional experiments using stopped-flow spectrophotometry were conducted to study the degradation of the metallocarbenoid intermediates in the presence of a suitable substrate. A solution containing a known concentration of ferrous C45/AP3.2/*Rma*-TDE (15 μM, 100 mM KCl, 20 mM CHES, pH 8.6, reduced with Na_2_S_2_O_4_) was placed in one syringe and an 80:20% ethanol:water solution containing 1 mM of a carbene precursor (either EDA, ^*t*^BuDA or BnDA) and 6 mM of styrene was placed in the second syringe. 50 μL from each syringe was simultaneously injected into a mixing chamber and the progression of the reaction was monitored spectroscopically, at 5 °C and 25 °C, over the course of 180-1000 seconds to examine carbene transfer activity. The progress of the reaction was monitored at 428-433 nm and 417 nm respectively. Final concentrations were 7.5 μM reduced C45/AP3.2/*Rma*-TDE, 500 μM EDA/^*t*^BuDA/BnDA, 3 mM styrene (40% ethanol).

### Protein mass spectrometry for C45

The formation of the C45-metallocarbenoid intermediate was further examined using positive electron-spray-ionization (ESI) mass spectrometry (Waters Xevo G2-XS QTof). A 1 mL solution of 350 μM C45 in CHES buffer (100 mM KCl, 20 mM CHES, pH 8.6), in a 1.5 mL sealed-top vial containing a silicon septum, was flushed under scrubbed nitrogen inside an anaerobic glovebox ([O_2_] < 5 ppm; Belle Technology) before being sealed, removed from the glovebox and placed on ice for 10 minutes. The sample was then immediately loaded directly into the mass spectrometer, where an isocratic 100 mM ammonium acetate solution was employed as the mobile phase (0.25 mL.min^-1^). 20 μL injections were employed and the progression of the sample into the chamber was monitored spectroscopically at 280 nm. The mass spectrum contained peaks screened across a m/z range of 1500-2700. The dominant peaks were identified, and the charges of the fragments were calculated using the molecular mass of C45 (15172.5 Da).

A 1 mL solution of 350 μM C45 in CHES buffer (100 mM KCl, 20 mM CHES, pH 8.6), in 1.5 mL sealed-top vials, was then flushed under scrubbed nitrogen inside an anaerobic glovebox ([O_2_] < 5 ppm; Belle Technology) followed by the addition of 25 μL of Na_2_S_2_O_4_ (400 mM stock in CHES buffer). A separate vial containing the selected diazo compound (EDA, ^*t*^BuDA, BnDA, 400 mM stock in EtOH) was deoxygenated alongside the vial containing reduced C45. The vials were sealed, transported out of the glovebox and cooled in an ice bath for 10 minutes. Once cooled, 50 μL of the selected diazo compound was added *via* gastight syringe into the vial containing reduced C45 to initiate the reaction. The sample was kept on ice for 1 minute to allow for the formation of the metallocarbenoid intermediate before being directly loaded onto the mass spectrometer. Final reaction concentrations were 350 μM enzyme, 10 mM sodium dithionite, and 20 mM diazo compound. Identical conditions to the experiment with only C45, mentioned above, were employed for each carbene precursor studied.

### Carbene transfer chemistry

Unless stated otherwise, all assays were conducted under scrubbed nitrogen inside an anaerobic glovebox ([O_2_] < 5 ppm; Belle Technology). The assays were conducted inside 1.5 mL screw top vials sealed with a silicone-septum containing cap. All assays were conducted in CHES buffer (100 mM KCl, 20 mM CHES, pH 8.6) except for the assays conducted with Mb(H64V,V68A) which were performed in KPi buffer (100 mM potassium phosphate, pH 7).The final reaction volumes for all assays were 400 μL unless otherwise stated.

### Cyclopropanation assays

To 370 μL of a 10 μM C45/AP3.2/*Rma*-TDE /Mb(H64V,V68A) solution was added 10 μL of Na_2_S_2_O_4_ (400 mM stock; de-ionized water) and the mixture was left to stir for 1 minute, ensuring complete reduction of C45 from Fe^3+^ to Fe^2+^. 10 μL of the selected styrene (1.2 M stock in EtOH) was added and the reaction left to mix for 30 seconds. A separate vial containing the selected diazo compound (400 mM stock in EtOH) was deoxygenated alongside the reaction vessel. The vials were sealed, transported out of the glovebox and cooled in an ice bath. Once cooled, 10 μL of the diazo compound was added *via* gastight syringe into the reaction vials to initiate the reaction. Final reaction concentrations were 10 μM enzyme (0.1% mol%), 10 mM sodium dithionite, 10 mM diazo compound, and 30 mM styrene. Once mixed, the reactions were stirred on a roller at room temperature. After 2 hours, the reaction was quenched by the addition of 20 μL of 3 M HCl. The vials were subsequently unscrewed, and 1 mL of ethyl acetate was added to the vial. The solution was transferred to a 1.5 mL microcentrifuge tube, vortexed and centrifuged for 1 minute at 13,500 rpm. The upper organic layer was extracted, dried over MgSO_4_ if necessary, and subsequently used for analysis. Where base hydrolysis of the resultant ester was necessary, 400 μL of 3 M NaOH was introduced to the organic layer and the mixture was left to stir, at room temperature for 30 minutes (the progress of the ester hydrolysis was monitored using TLC (7:3 ethyl acetate:hexane)). Products were analyzed by chiral-HPLC and LC-MS as described below. The product yields, enantiomeric excesses and total turnover numbers (TTN; concentration of product formed/concentration of enzyme) were calculated *via* external calibration with commercial ethyl 2-phenylcyclopropane-1-carboxylate and 2-phenylcyclopropane-1-carboxylic acid.

### Carbonyl olefination assays

Carbonyl olefination assays were conducted under the same reaction conditions as the cyclopropanation assays, with the following alterations: alkene substrates were substituted for selected benzaldehyde substrates; 10 mM of PPh_3_ was included as an additional reagent in the reaction (10 μL of a 400 mM stock in acetone); the reactions were carried out overnight; after quenching, the products were extracted with 1 mL of dichloromethane; the products were analyzed by C18 reverse-phase HPLC and LC-MS as described below. The product yields, cis/trans ratios and total turnover numbers where calculated *via* an external calibration with commercial ethyl cinnamate and cinnamic acid.

### N-H insertion assays

N-H insertion assays were conducted under the same reaction conditions as the cyclopropanation assays, with the following alterations: the alkene starting materials were substituted for the selected amines; the reactions were carried out overnight; after quenching, the products were extracted with 1.25 mL of *n*-hexane (1.25 mL of hexane as required for sufficient extraction of the product into the organic phase [monitored by TLC]); the products were subsequently analyzed by chiral-HPLC and LC-MS as described below. The product yield and total turnover numbers where calculated *via* an external calibration with commercial *n*-phenyglycine ethyl ester and *n*-ethylpiperidine acetate (and including a 1.25 multiplication factor to account for product dilutions).

### Ring expansion assays

Ring expansion assays were conducted under similar reaction conditions as the cyclopropanation assays described above, but with the following alterations: styrene was substituted for pyrrole (400 mM stock in ethanol); ethyl 2-bromo-2-diazoacetate was used as the carbene precursor (40 mM stock in CH_2_Cl_2_); the products were extracted with 1 mL of ethyl acetate and subsequently analyzed by C18 reverse-phase HPLC and LC-MS as described below. Final reaction concentrations were 10 μM enzyme (1% mol%), 10 mM sodium dithionite, 1 mM diazo compound, and 10 mM pyrrole. The product yields and total turnover numbers where calculated *via* an external calibration with commercial ethyl nicotinate and niacin.

### Product characterization by reverse phase and chiral HPLC

All the reactions performed using C45 and AP3.2 were quantified by High Performance Liquid Chromatography (HPLC). Two separate columns were employed for quantification: a chiral column and a reverse phase C18 column.

A chiral-HPLC column (Astec CHIROBIOTIC^®^ V, 250 x 21 mm, 5 μm) was used to analytically quantify the cyclopropanation and N-H insertion assays, employing an isocratic polar organic mobile phase (100% CH_3_CN: 0.1% v/v TFA: 0.1% v/v: Et_3_N; 0.1 mL.min^-1^ flow rate and 2 μL injection volume for the cyclopropanation assays; 0.2 mL.min^-1^ flow rate and 5 μL injection volume for the N-H insertion assays). Injection volumes of the N-H insertion assay products were 5 μL to account for the 2.5:1 dilution in the extraction step. All elution traces were monitored spectroscopically at 245, 254 and 280 nm. The chiral-HPLC column allowed the retention times of the starting materials and the reaction products to be determined, and to quantify, where necessary, the enantioselectivity of a given reaction.

A C18 HPLC reverse phase column (Phenomenex, 150 x 15 mm, 5 μm) was used to analytically quantify the ring expansion and carbonyl olefination assays, employing a gradient as the mobile phase for ring expansion assays (70:30% H_2_O:CH_3_CN to 10:90% H_2_O:CH_3_CN; 2 mL.min^-1^ flow rate and 20 μL injection volume) and an isocratic mobile phase for the carbonyl olefination assays (100% CH_3_CN: 0.1% v/v TFA: 0.1% v/v: Et_3_N; 2 mL.min^-1^ flow rate and 20 μL injection volume). Injection volumes of the sample mixture were 20 μL to account for the 2.5:1 dilution in the extraction step. All elution traces were monitored spectroscopically at 245, 254, 265 and 280 nm. The C18 column allowed the retention times of the starting materials and the reaction products to be determined, and, where necessary, to quantify the diastereoselectivity of a given reaction.

If appreciable product formation could be detected after the initial HPLC experiments, the products were subsequently analyzed using Liquid chromatography-Mass spectrometry (LC-MS) to assist in product identification (described below).

External calibrations of the anticipated reaction products were conducted using HPLC with commercially available standards. The conditions and mobile phases employed in the product analysis for each assay was employed for the calibrations. Each substrate was prepared at multiple concentrations (i.e. 50-500 μM, 1 mM, 2 mM, 5 mM, 7.5 mM, 10 mM, and 20 mM) and the peak height response in the chromatogram was recorded as a function of concentration. A plot of [substrate] vs peak height engendered a straight line which could be used to determine the product yields and TTNs for each assay. Injection volumes for the external calibrations were 2 μL and 8 μL for the chiral-HPLC and C18 columns respectively.

### Product characterization by liquid chromatography-mass spectrometry

Liquid chromatography-Mass spectrometry (LC-MS, C8) was employed to identify the products formed in each assay. A C8 reverse column (Grace Vydac, 100 x 21 mm, 5 μm) was used with a 10 minute gradient mobile phase (95:5% H_2_O:CH_3_CN to 10:90% H_2_O:CH_3_CN; 0.1% v/v formic acid, 0.25 mL.min^-1^). Injection volumes of 20 μL were employed and the chromatogram was screened across a wavelength range of 240-300 nm. After eluting from the column, the mixture entered an isocratic solvent chamber where a 1:100 dilution was performed prior to injecting the sample into a positive electron-spray-ionization (ESI) mass spectrometer (Waters Xevo G2-XS QTof). The mass spectrum contained peaks screened across a m/z range of 70-250. Retention times and fragmentation patterns were initially determined using commercial samples of the anticipated products.

### Circular dichroism spectroscopy

Circular Dichroism spectra were collected using a JASCO J-815 CD polarimeter. AP3.2 was loaded into a 1 mm pathlength quartz cell at 0.01 mg.mL^-1^ in mM KCl, 20 mM CHES, pH 8.58, and far-UV CD spectra were recorded at 100 nm/min with a sensitivity of 50 mdeg. Thermal stability of AP3.2 was assessed by monitoring ellipticity at 222 nm while increasing temperature at a ramp rate of 40 °C/hour with 1 °C intervals. All raw data was converted to mean residue ellipticity (MRE) and the thermal denaturation transition midpoint (*T_m_*) was assessed by plotting the second derivatives of a smoothed thermal denaturation trace where the x-axis intercept corresponds to the *T_m_*.

### Redox potentiometry

The redox potential of AP3.2 was measured as previously described for C45^6^. Briefly, AP3.2 (50 μM) was loaded into a spectroelectrochemical cell in 100 mM KCl, 50 mM CHES, pH 8.6, 10% glycerol with the following redox mediators: 20 μM benzyl viologen, 20 μM anthroquinone-2-sulfonate, 20 μM phenazine, 25 μM 2-hydroxy-1,4-napthoquinone. Potential was applied to the spectroelectrochemical cell (platinum working and counter electrodes, Ag/AgCl reference electrode) using a Biologic SP-150 potentiostat in both reductive and oxidative directions to confirm equilibration. AP3.2 heme reduction potential was calculated by plotting the fraction of reduced protein versus the applied potential, and the data were fitted using the following single electron Nernst function:

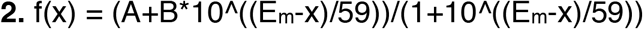

In equation **2**, A and B are y-axis values at 100% oxidized and reduced heme respectively; Em is the heme reduction potential.

### Whole cell C45-catalysed ring expansion experiments

Overnight starter cultures were prepared by adding 100 μL carbenicillin (50 mg.mL^-1^) and 100 μL chloramphenicol (50 mg.mL^-1^, C45 only) to 100 mL of LB before inoculating the media with a C45-expressing *E. coli* glycerol stock. Starter cultures were then incubated overnight at 37 °C and 180 rpm. 50 mL of the overnight starter culture was then used to inoculate 1 L of LB containing the same concentrations of antibiotics (see above). The 1 L cultures were grown in a shaking incubator at 37 °C and 180 rpm until an OD_600_ between 0.6-0.8 was obtained (usually after 3 hours), at which point 980 μL was extracted from both vessels, placed inside separate 1.5 mL screw top vials and placed on ice. 1 mL of IPTG solution (1 M stock, 1 mM final concentration at induction) was added to specific cultures to induce protein expression; induced and non-induced cultures were left in the incubator for an additional three hours (37 °C and 180 rpm). After three hours, 980 μL was extracted from the vessels and transferred to separate 1.5 mL screw top vials. The samples were then degassed inside an anaerobic glove-box (Belle Technology) for 30 minutes before 10 μL of pyrrole (1 M stock in EtOH) was added and the vials sealed and removed from the glovebox. The vials were cooled on ice before 25 μL of ethyl 2-bromo-2-diazoacetate (40 mM stock in DCM, nitrogen flushed) was added *via* a gastight needle; the final concentration of the reagents were 1 mM ethyl 2-bromo-2-diazoacetate and 10 mM pyrrole. After the reactions had been left stirring for 2 hours, the samples were quenched with 3 M HCl (30 μL) and extracted with 1 mL ethyl acetate. Analysis was conducted as reported above in the “Ring expansion assays’’ section.

### Hydrolysis of ethyl nicotinate by a *Bacillus subtilis* esterase

To a 100 mL round-bottom flask equipped with a small magnetic stirrer bar was added 19.8 mL of CHES buffer (pH 8.6) and 200 μL of commercial ethyl nicotinate (5 M stock in DMSO). The final concentration of ethyl nicotinate was 50 mM. A 100 μL aliquot of the solution was extracted to an Eppendorf tube for HPLC analysis. 2 mg of esterase from *Bacillus subtilis* (SigmaAldrich) was added to the mixture, then the round-bottom flask was closed with a stopper and the solution was left to stir for 1 hour. After 1 hour, a 1 mL aliquot of the solution was extracted into a 1.5 mL Eppendorf tube, and 100 μL of 3 M trichloroacetate was added to precipitate the protein. The mixture was vortexed, centrifuged (30 seconds, 13,000 rpm) and the resulting solution analyzed directly *via* HPLC. The HPLC protocol for the analysis was identical to the procedure outlined in the ‘’ring expansion assays’’ section above.

## Acknowledgements

This work was supported at the University of Bristol by the BBSRC (grant no: BBI014063/1, BB/R016445/1 & BB/M025624/1), the Bristol Centre for Functional Nanomaterials (EPSRC Doctoral Training Centre Grant EP/G036780/1) through a studentship for R.S. and by the SynBioCDT (EPSRC and BBSRC Centre for Doctoral training in Synthetic Biology EP/L016494/1) through a studentship for J.W.S. We also wish to thank Dr. Peter Wilson for his assistance in collecting LC-MS and MS data, and Dr Steve Burston for helpful kinetic discussions.

## Data Availability

The data that support the findings of this study are available from the corresponding author upon reasonable request.

## Supporting Information

Spectral and kinetic data, HPLC and chiral HPLC chromatograms, LC-MS and MS data, HPLC calibrations, circular dichroism spectra and thermal melts, a computational protein model mapped with AP3.2 mutations and a scheme of NAD biosynthesis are provided.

