## Supplementary information for "A *de novo* peroxidase is also a promiscuous yet stereoselective carbene transferase"

### Supporting Information

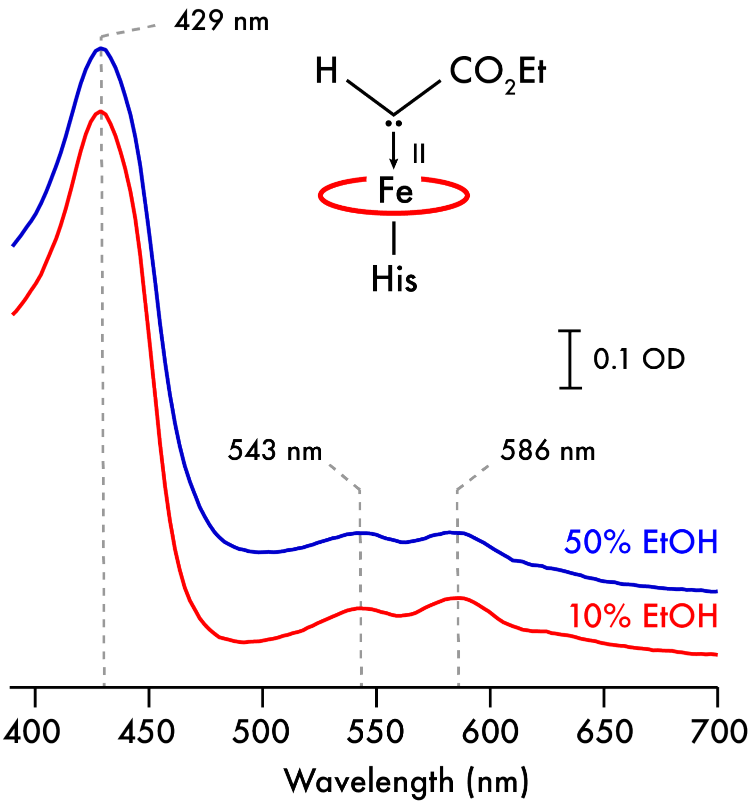


**Figure S1. The effect of ethanol on the metallocarbenoid spectra of C45.** Electronic spectra were recorded after rapid mixing of ferrous C45 (7.5 µM) with EDA (500 µM) in 10 and 50% ethanol:buffer solutions at 5 °C.

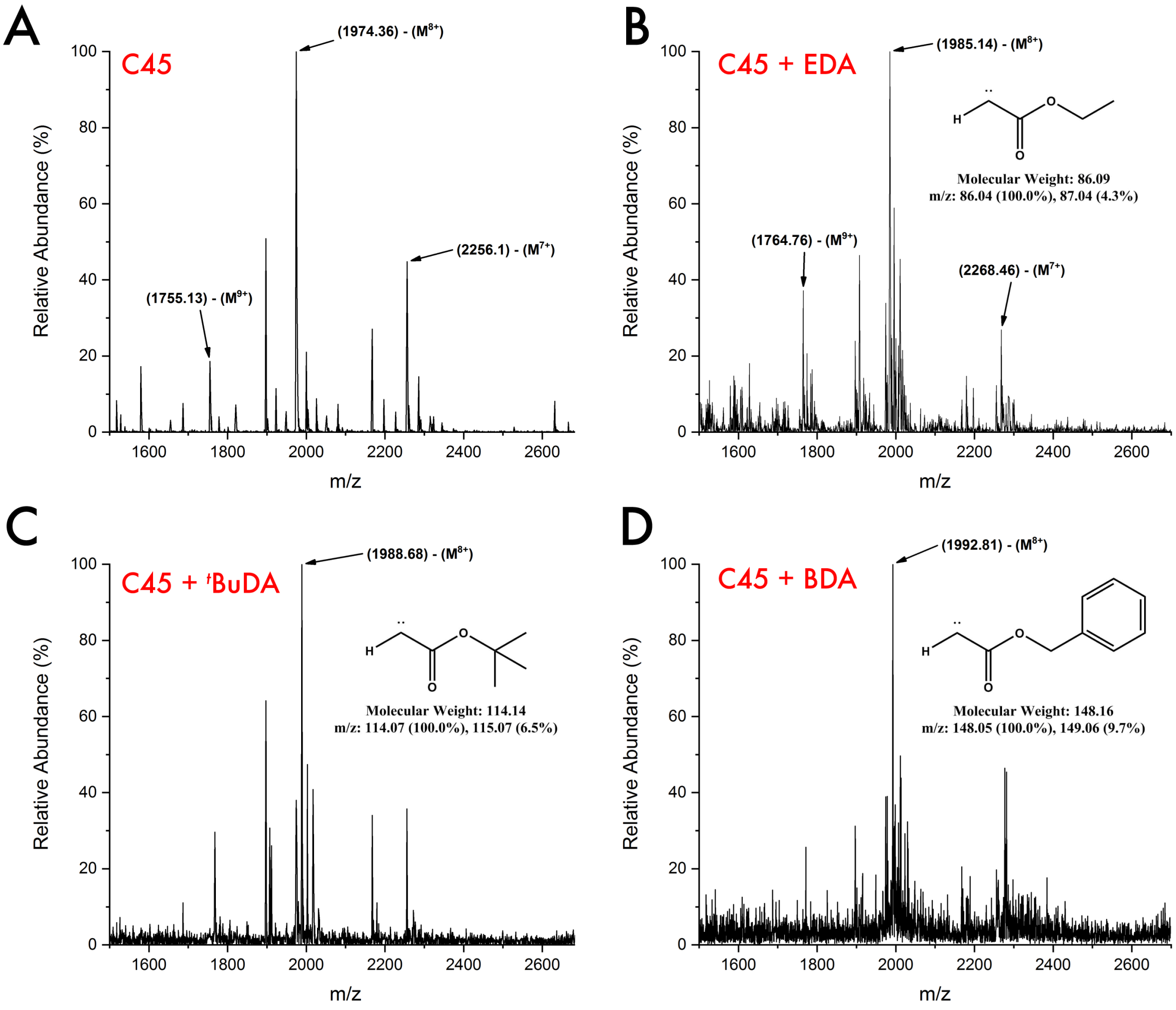


**Figure S2. ESI mass spectrometry of C45 metallocarbenoid complexes. A.** Mass spectrum of C45 (350 µM) in buffer. The three dominant peaks occur at m/z values of 1755.13, 1974.36, and 2256.1 Da respectively. These fragments correspond to a charged species of (from left to right) $9^{+}$, $8^{+}$, and $7^{+}$, as determined by the atomic mass of C45 (15793.17 Da). **B.** Mass spectrum for C45 after the addition of 50 µl of ethyl diazoacetate (EDA, 20 mM, 5% ethanol). The three dominant peaks in the spectrum have shifted, relative to the peaks exhibited in the C45 spectrum, to 1764.76, 1985.14, and 2268.46 Da for the $M^{9+}$, $M^{8+}$ and $M^{7+}$ species respectively. The spectrum indicates an average mass increase of 86.52 Da, which corresponds approximately with the predicated molecular weight of an ethyl diazoacetate carbene ($-N_2$) species (86.09 Da). **C.** Mass spectrum for C45 after the addition of 50 µl of *tert*-butyl diazoacetate ($^{t}$BuDA, 20 mM, 5% ethanol). The dominant peak corresponding to the $M^{8+}$ species has shifted to an m/z value 1988.68, indicating an increase in mass of 114.56 Da, which corresponds with the predicated molecular weight of a *tert*-butyl diazoacetate carbene ($-N_2$) species (114.14 Da). **D.** Mass spectrum for C45 after the addition of 50 µl of benzyl diazoacetate (BnDA, 20 mM, 5% ethanol). The dominant peak corresponding to the $M^{8+}$ species has shifted to an m/z value 1992.81, indicating an increase in mass of 148.08 Da, which corresponds with the predicated molecular with of a benzyl diazoacetate carbene ($-N_2$) species (148.05 Da).

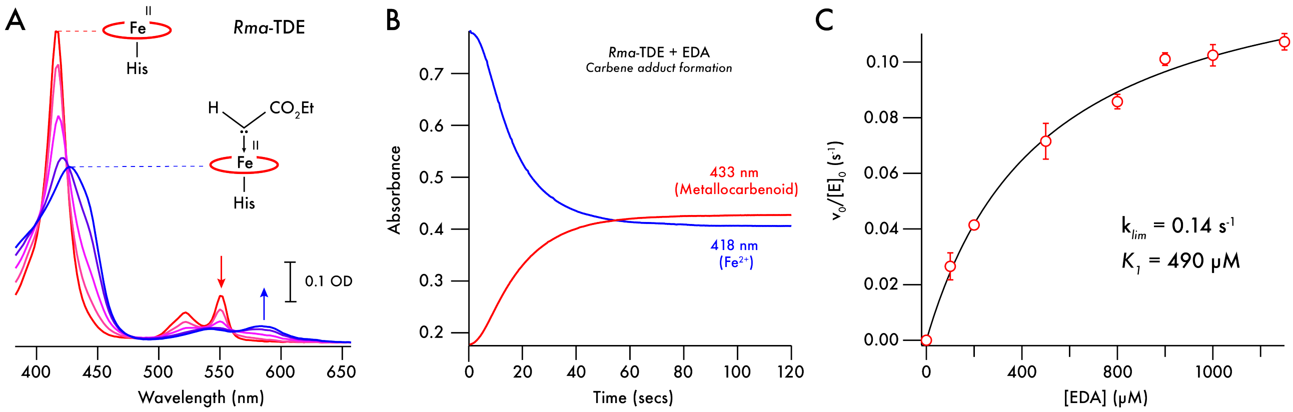


**Figure S3. Metallocarbenoid formation and kinetics of *Rma*-TDE with EDA. A.** Time course of electronic spectra recorded following rapid mixing of ferrous *Rma*-TDE (7.5 μM, red spectrum) with EDA in 40% EtOH at 5 °C. The appearance of the metallocarbenoid intermediate (blue spectrum) is concomitant with the disappearance of the ferrous *Rma*-TDE spectrum. Spectra presented were recorded 1, 2, 4, 6, 8, 10, 15, 20, 50 and 120 seconds after mixing. **B.** Metallocarbenoid formation and stability in the absence of styrene substrate. Single wavelength traces represent the time course of ferrous *Rma*-TDE (418 nm, blue; 7.5 μM protein, 100 μM EDA, 40% EtOH, 20 mM CHES, 100 mM KCl, pH 8.6) and metallocarbenoid:*Rma*-TDE adduct (433 nm, red) following rapid mixing of ferrous *Rma*-TDE with 500 μM ethyl diazoacetate at 5 °C. **C.** EDA-concentration-dependent formation of the *Rma*-TDE metallocarbenoid adduct. Kinetic data were recorded using a stopped-flow spectrophotometer and analyzed as described in the Materials and Methods. The limiting rate constant ($k_{lim}$) and *pseudo*-Michaelis constant ($K_1$) for metallocarbenoid formation are 0.14 $s^{-1}$ and 490 μM respectively.

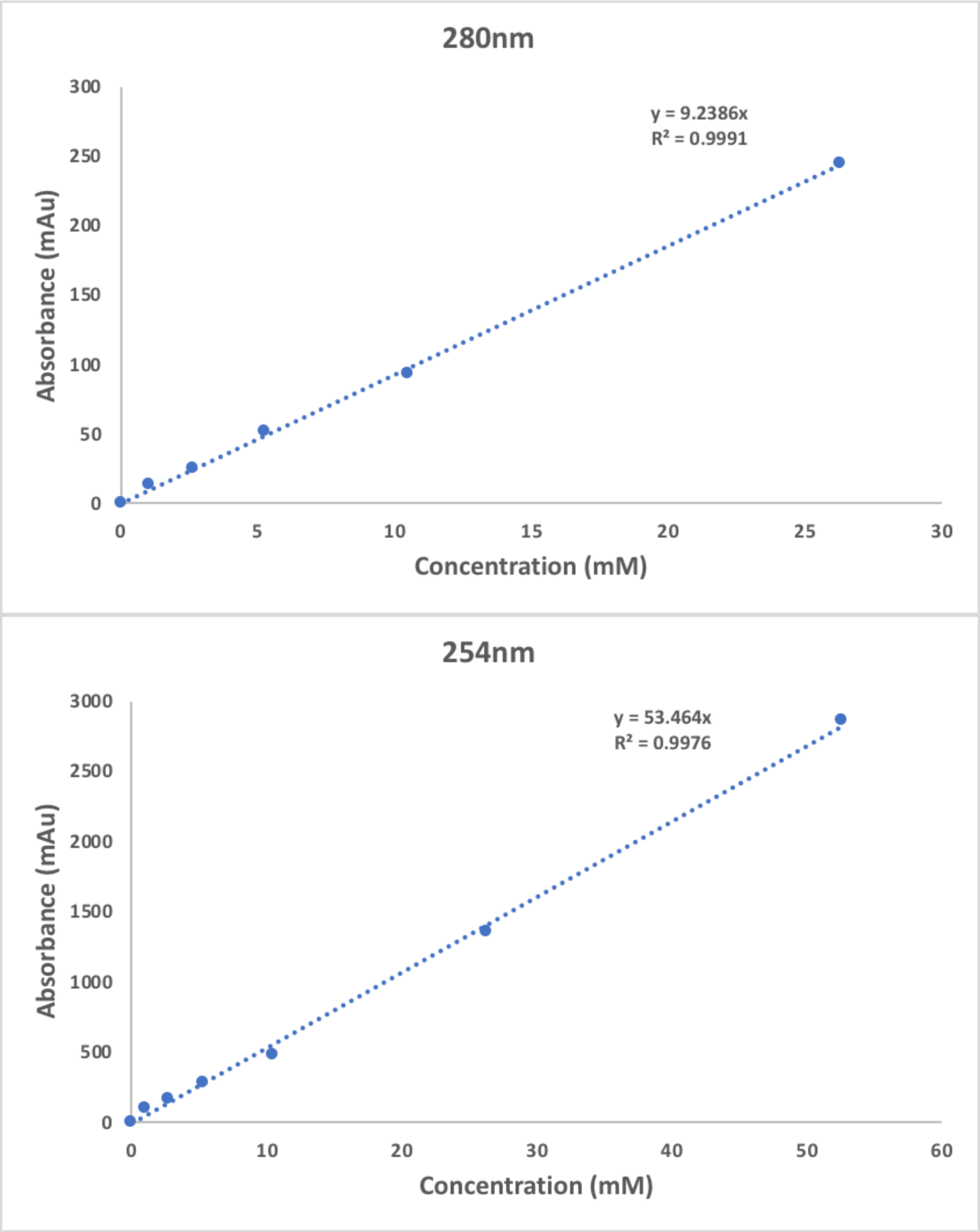


**Figure S4. Chiral-HPLC external calibrations for ethyl 2-phenylcyclopropane-1-carboxylate at 280 nm (upper panel) and 254 nm (lower panel).** A polar organic mobile phase (100% MeCN: 0.1% v/v TFA: 0.1% v/v: $Et_3N$) was employed and injection volumes were 2 µl.

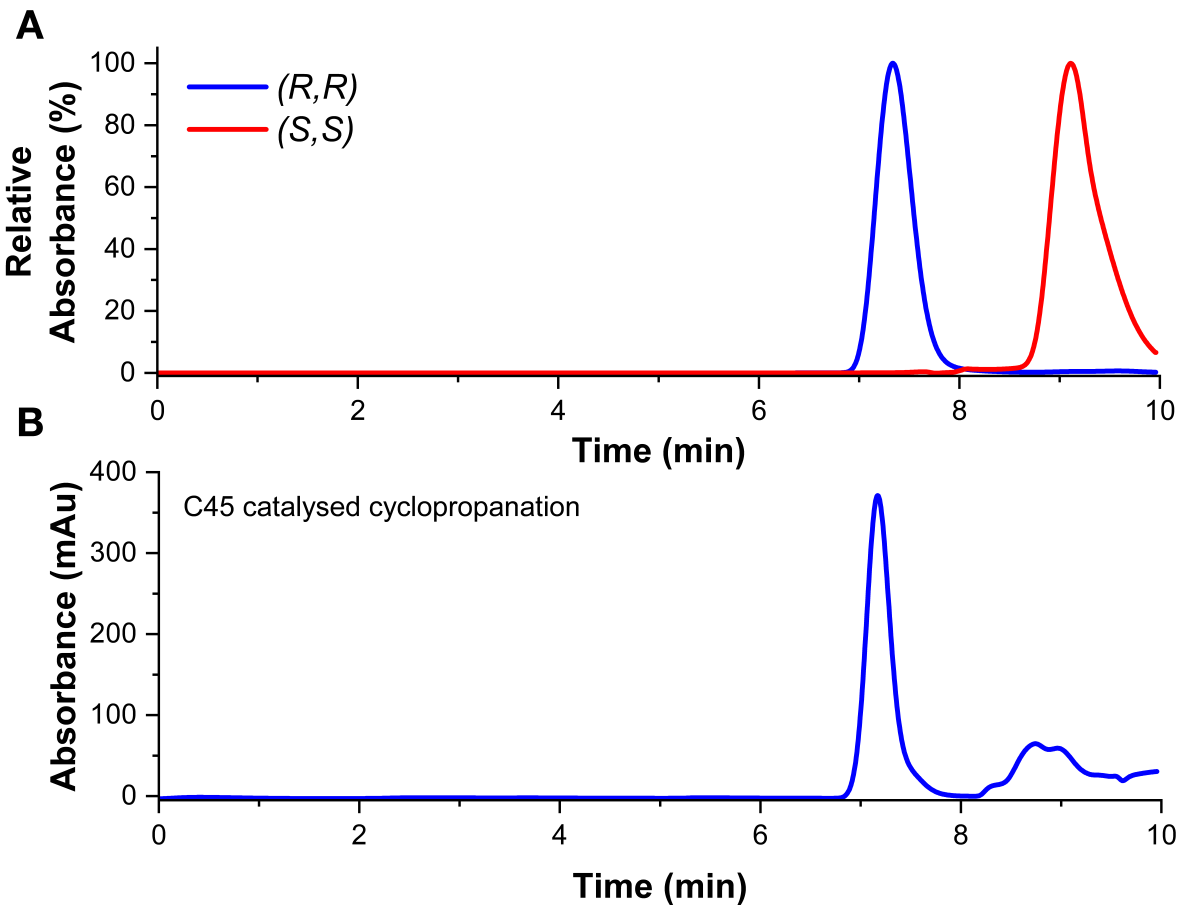


**Figure S5**. **Chiral-HPLC chromatograms for the cyclopropanation assays.** A polar organic mobile phase (100% MeCN: 0.1% v/v TFA:0.1% v/v: $Et_3N$) was employed and injection volumes were 2 µl. **A.** Normalised commercial (*R,R*)-ethyl 2-phenylcyclopropane-1-carboxylate (254 nm, EtOH, blue) and (S,S)-ethyl 2-phenylcyclopropane-1-carboxylate (254 nm, EtOH, red). **B.** Averaged chromatogram from the C45 (10 µM, 0.1% catalyst loading) catalyzed cyclopropanation assay between styrene (30 mM) and EDA (10 mM) (CHES buffer, pH 8.6, 254 nm). The cyclopropane product from each assay was extracted with 1 ml of ethyl acetate and 400 µl of 3M NaOH prior to loading onto the column. The (R,R)-enantiomer eluted first and was followed by the (S,S)-enantiomer. The relative peak heights for the (R,R) and (S,S) enantiomers was used to calculate enantiomeric excess values using the equation ([R,R]-[S,S])/ ([R,R]+ [S,S]).

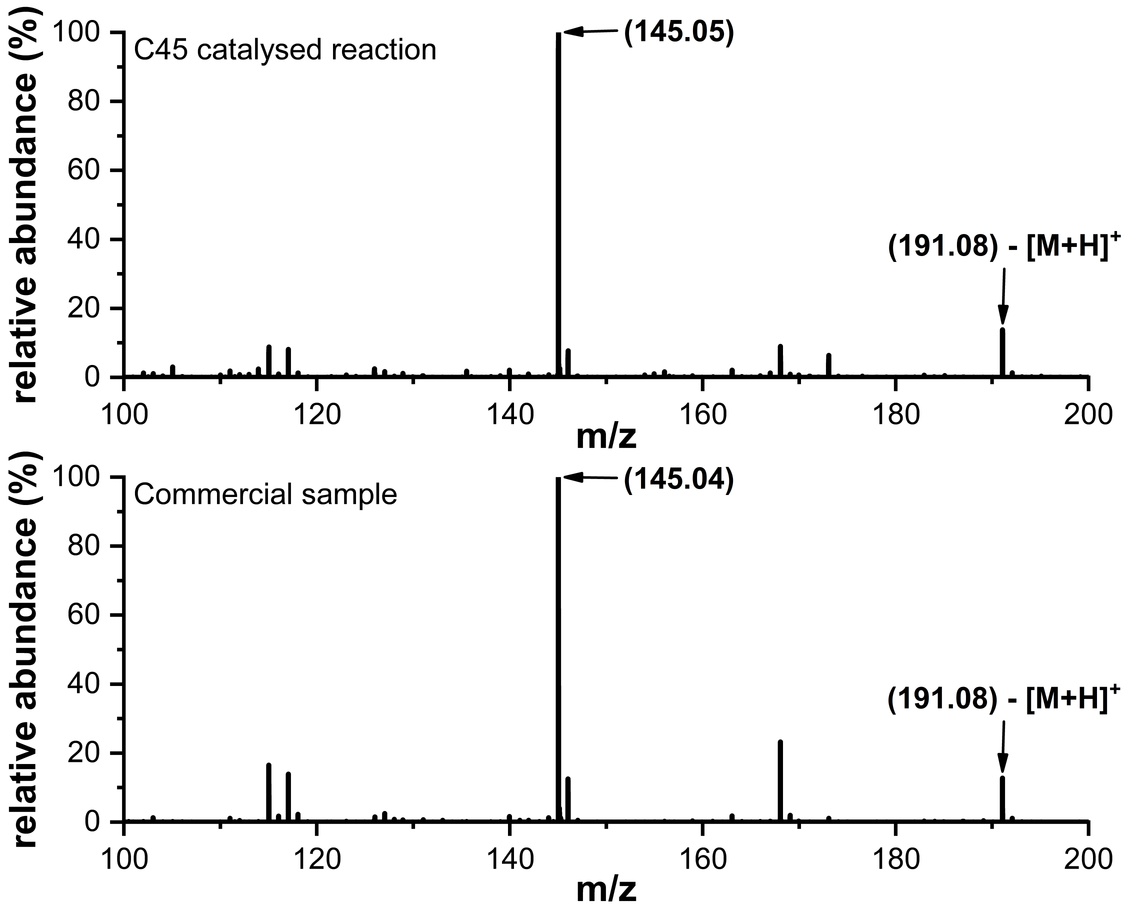


**Figure S6. LC-MS spectra of C45 catalyzed cyclopropanation assay products.** Commercial ethyl 2-phenylcyclopropane-1-carboxylate (in EtOH) exhibiting the dominant oxonium ion fragment at 145 m/z (top MS) and C45 catalyzed cyclopropanation assay between styrene (30 mM) and EDA (10 mM) (bottom MS). All spectra were recorded in ES+ mode and monitored at 254 and 280 nm. A C8 column was employed for the LC separation with a gradient mobile phase (95:5:0.1% v/v water/MeCN/formate 10:90:0.1% v/v water/MeCN/formate). Assignment of major product peaks in the mass spectra can be found in SI Fig. S9, where R = H.

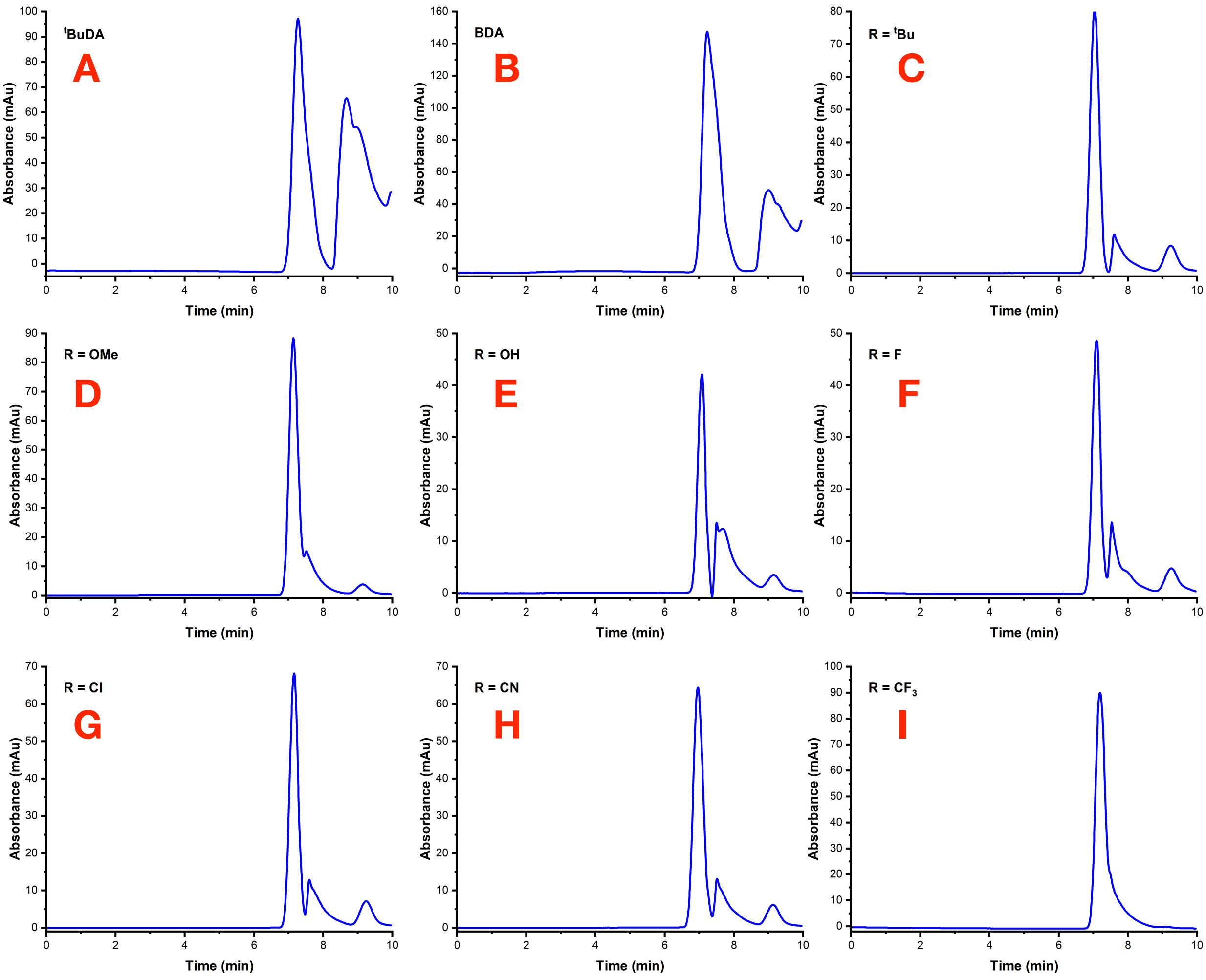


**Figure S7. Chiral-HPLC chromatogram for the C45 (10 µM) catalyzed cyclopropanation assays. A.** styrene (30 mM) and *tert*-butyl diazoacetate (10 mM), **B.** styrene (30 mM) and benzyl diazoacetate (10 mM), **C.** *p*-*tert*-butylstyrene (30 mM) and EDA (10 mM), **D.** *p*-methoxystyrene (30 mM) and EDA (10 mM), **E.** *p*-hydroxystyrene (30 mM) and EDA (10 mM), **F.** *p*-fluorostyrene (30 mM) and EDA (10 mM), **G.** *p*-chlorostyrene (30 mM) and EDA (10 mM), **H.** *p*-cyanostyrene (30 mM) and EDA (10 mM), and **I.** *p*-trifluoromethylstyrene (30mM) and EDA (10 mM) . All assays were performed in CHES buffer (pH 8.6) with 10 µM C45 (0.1% catalyst loading). The cyclopropane product from each assay was extracted with 1 ml of ethyl acetate and 400 µl of 3 M NaOH prior to loading onto the column. A polar organic mobile phase (100% MeCN: 0.1% v/v TFA: 0.1% v/v: $Et_3N$) was employed and injection volumes were 2 µl; all traces were recorded at 280 nm. The (R,R)-enantiomer eluted first and was followed by the (S,S)-enantiomer. The relative peak heights for the (R,R) and (S,S) enantiomers was used to calculate enantiomeric excess values using the equation ([R,R]-[S,S])/ ([R,R]+[S,S]).

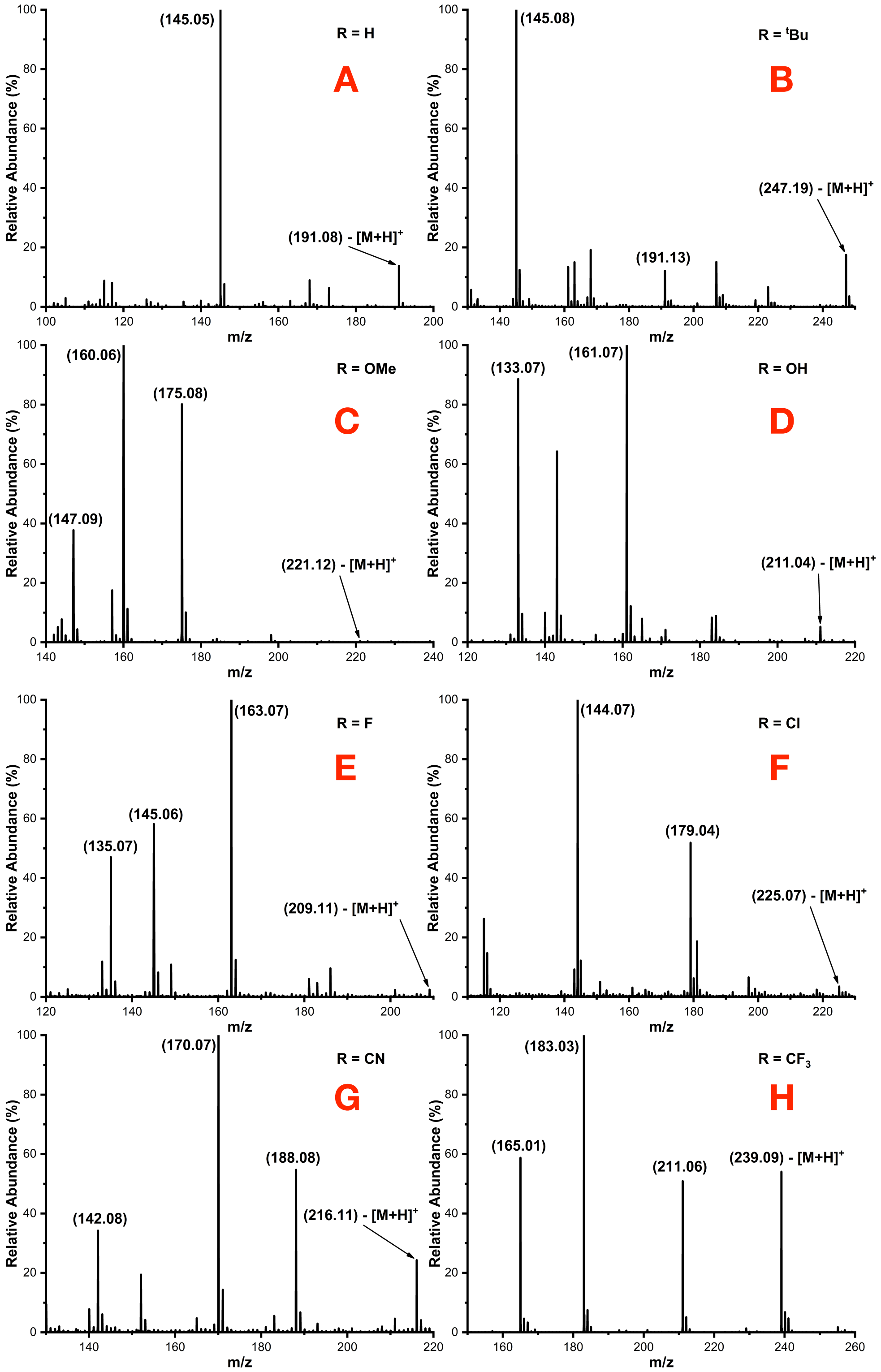


**Figure S8. LC-MS spectra and assigned fragments to major product peaks for the C45 (10 μM) catalyzed cyclopropanation assays (Top) A.** styrene **B.** *p-tert*-butylstyrene, **C.** *p*-methoxystyrene, **D.** *p*-hydroxystyrene, **E.** *p*-fluorostyrene, **F.** *p*-chlorostyrene, **G.** *p*-cyanostyrene, and **H.** *p*-trifluoromethylstyrene. All spectra were recorded in ES+ mode and monitored at 254 and 280 nm. A C8 column was employed for the LC separation with a gradient mobile phase (95:5:0.1% v/v water/MeCN/ formate 10:90:0.1% v/v water/MeCN/formate).

**A**

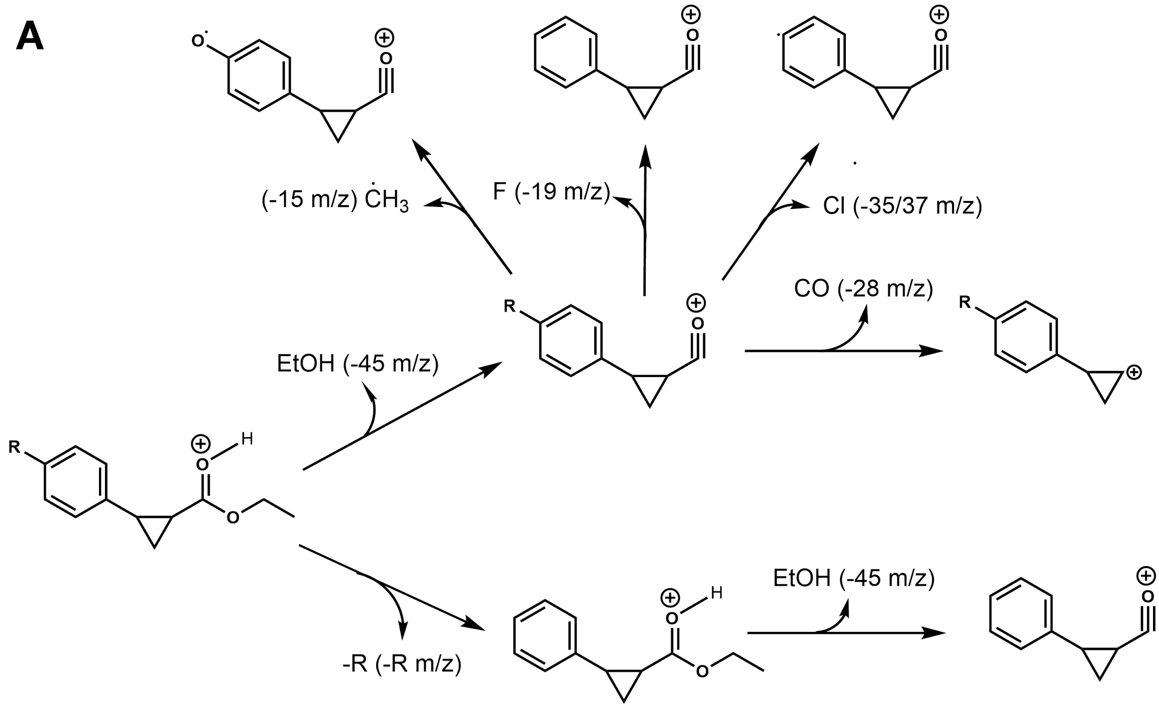


**B**

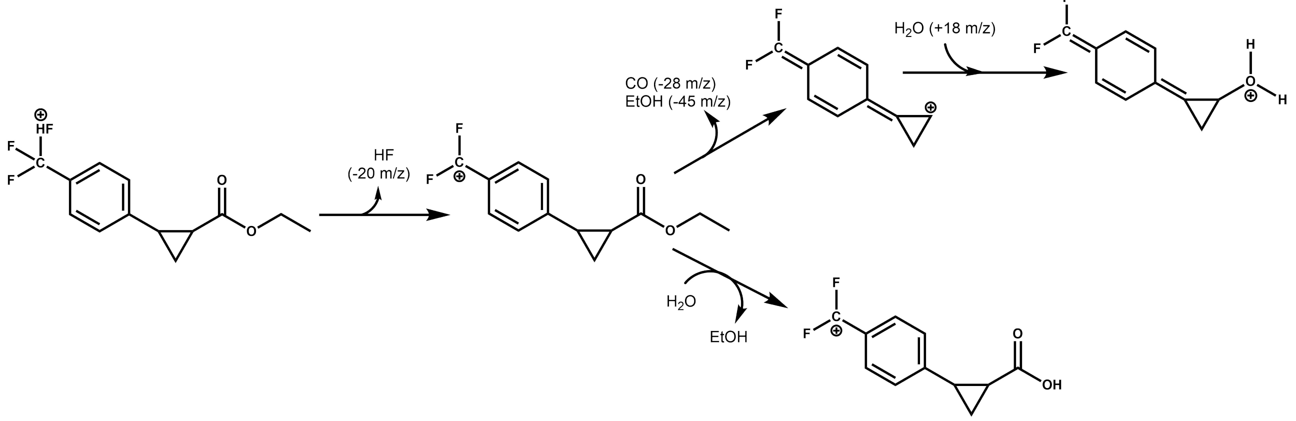


**Figure S9. Proposed fragmentation pathways for the products of C45-catalyzed cyclopropanation of *para*-substituted styrenes. A.** Fragmentation pathways leading to the observed peaks in the mass spectra. **B.** Fragmentation pathway for the product obtained using *p*-trifluoromethylstyrene as the starting substrate, resulting in the peaks observed in the product mass spectrum.

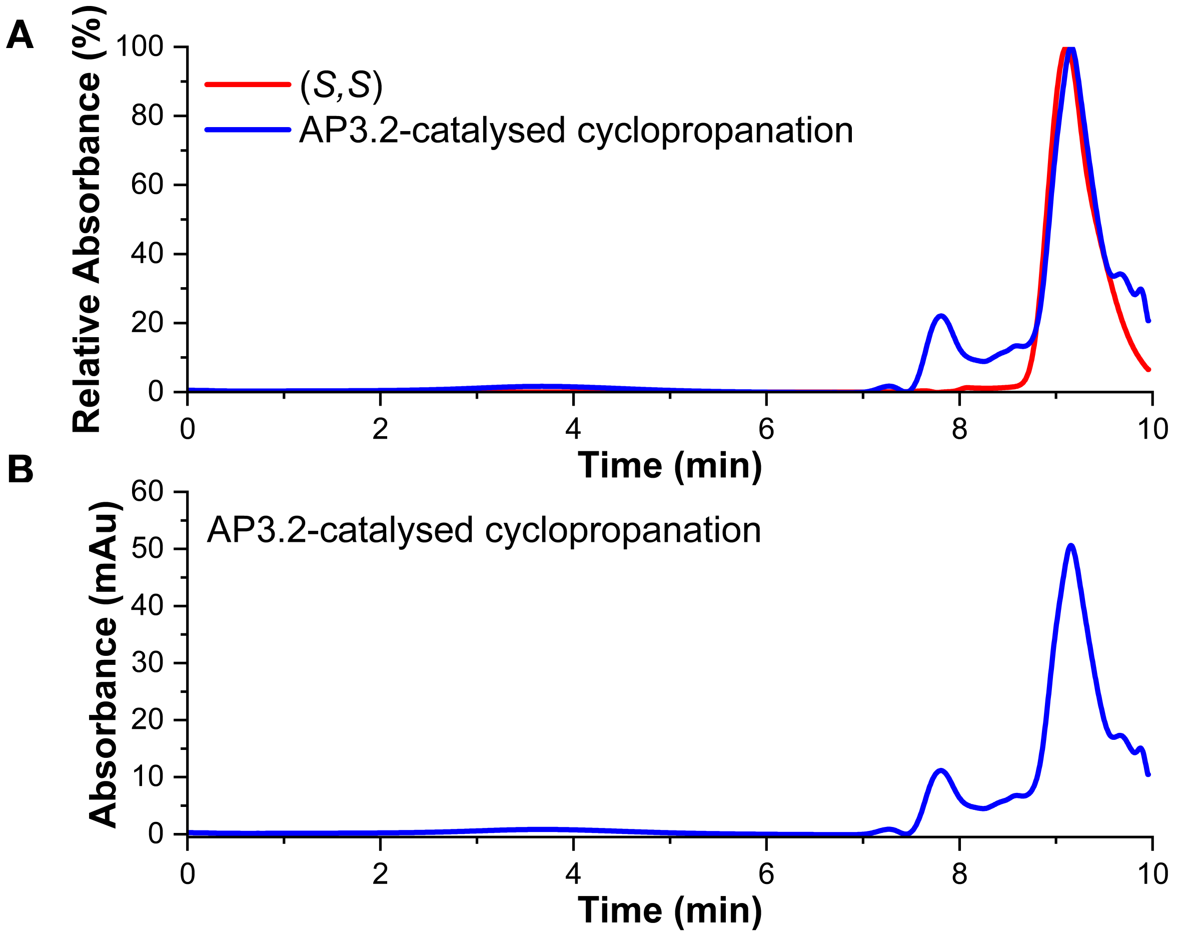


**Figure S10**. **Chiral-HPLC chromatograms for the AP3.2 catalyzed cyclopropanation assays. A.** normalised AP3.2 (10 µM, 0.1% catalyst loading) catalyzed cyclopropanation assay between styrene (30 mM) and EDA (10 mM) (100 mM KCl, 20 mM CHES, pH 8.6, EtOH, 254 and 280 nm) (blue line) vs normalised commercial (S,S)-ethyl 2-phenylcyclopropane-1-carboxylate (red line). **B.** Averaged AP3.2 (10 µM) catalyzed cyclopropanation assay between styrene (30 mM) and EDA (10 mM) (CHES buffer, pH 8.6, 254 and 280 nm). The cyclopropane product from each assay was extracted with 1 ml of ethyl acetate and 400 µl of 3M NaOH prior to loading onto the column. A polar organic mobile phase (100% MeCN: 0.1% v/v TFA:0.1% v/v: $Et_3N$) was employed and injection volumes were 2 µl. The relative peak heights for the (R,R) and (S,S) enantiomers was used to calculate enantiomeric excess values using the equation ([R,R]-[S,S])/ ([R,R]+[S,S]).

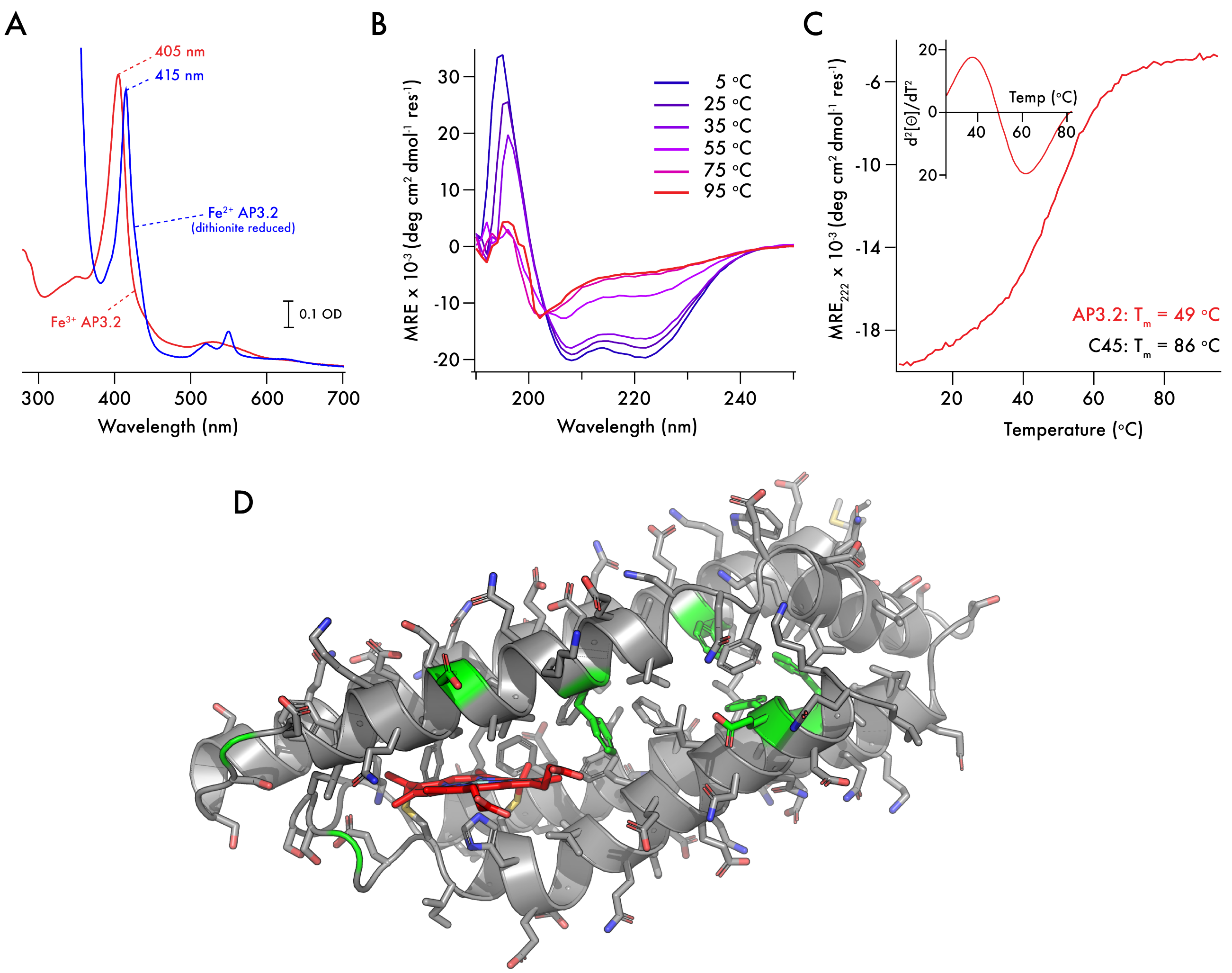


**Figure S11. Characterisation of AP3.2. A.** UV/visible spectra of ferric (red) and dithionite reduced ferrous AP3.2 (blue). **B.** Far-UV circular dichroism spectra of AP3.2 with varying temperature collected in 100 mM KCl, 20 mM CHES, pH 8.6. **C.** Temperature dependence of the CD signal monitored at 222 nm during thermal denaturation. The inset shows a smoothed second derivative of the thermal melt trace indicating a melting transition ($T_m$) of 49 °C. **D.** The positions of the eight mutations (F11Y/G39S/D48Y/F53S/F83S/G109A/F132S/E133G) are indicated in green on a computationally-derived model of C45[6].

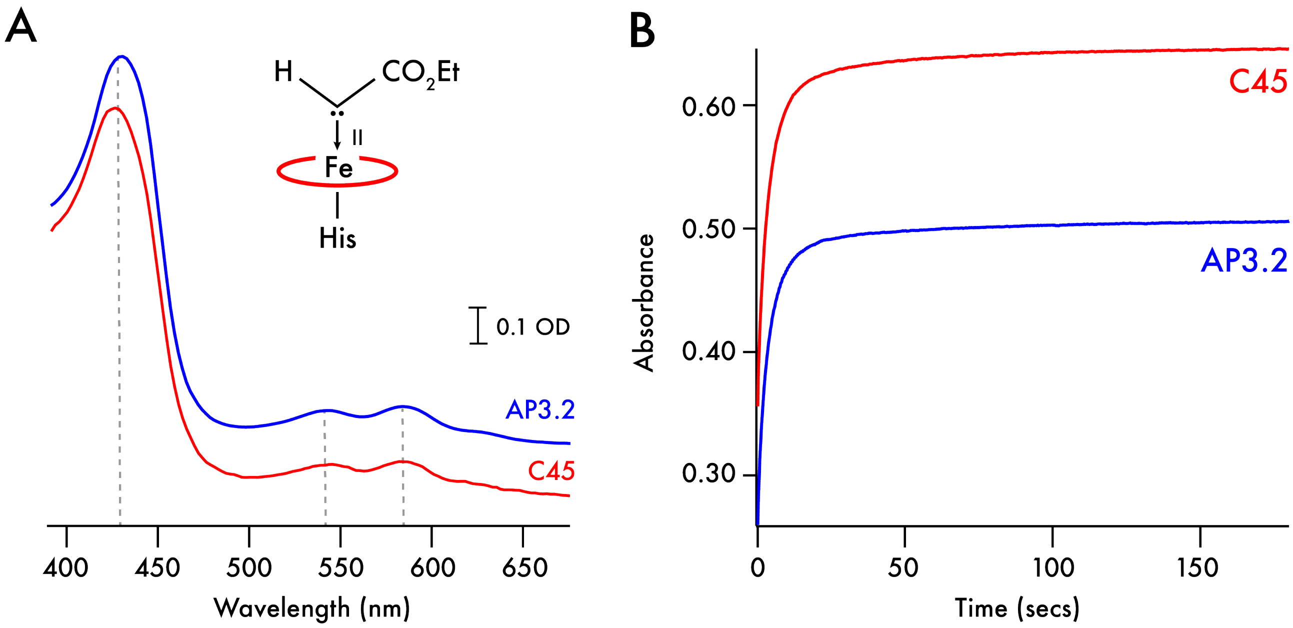


**Figure S12. Comparison of metallocarbenoid formation in C45 and AP3.2. A.** Visible spectra of the metallocarbenoid intermediates observed for C45 (red) and AP3.2 (blue) after rapid mixing of maquette (7.5 µM final concentration) with 2.5 mM EDA (40% EtOH) at 5 °C in the stopped-flow. **B.** Kinetic traces of metallocarbenoid formation in AP3.2 (blue) and C45 (red) measured at 437 nm after rapid mixing of maquette with 2.5 mM EDA (40% EtOH) at 5 °C in the stopped flow.

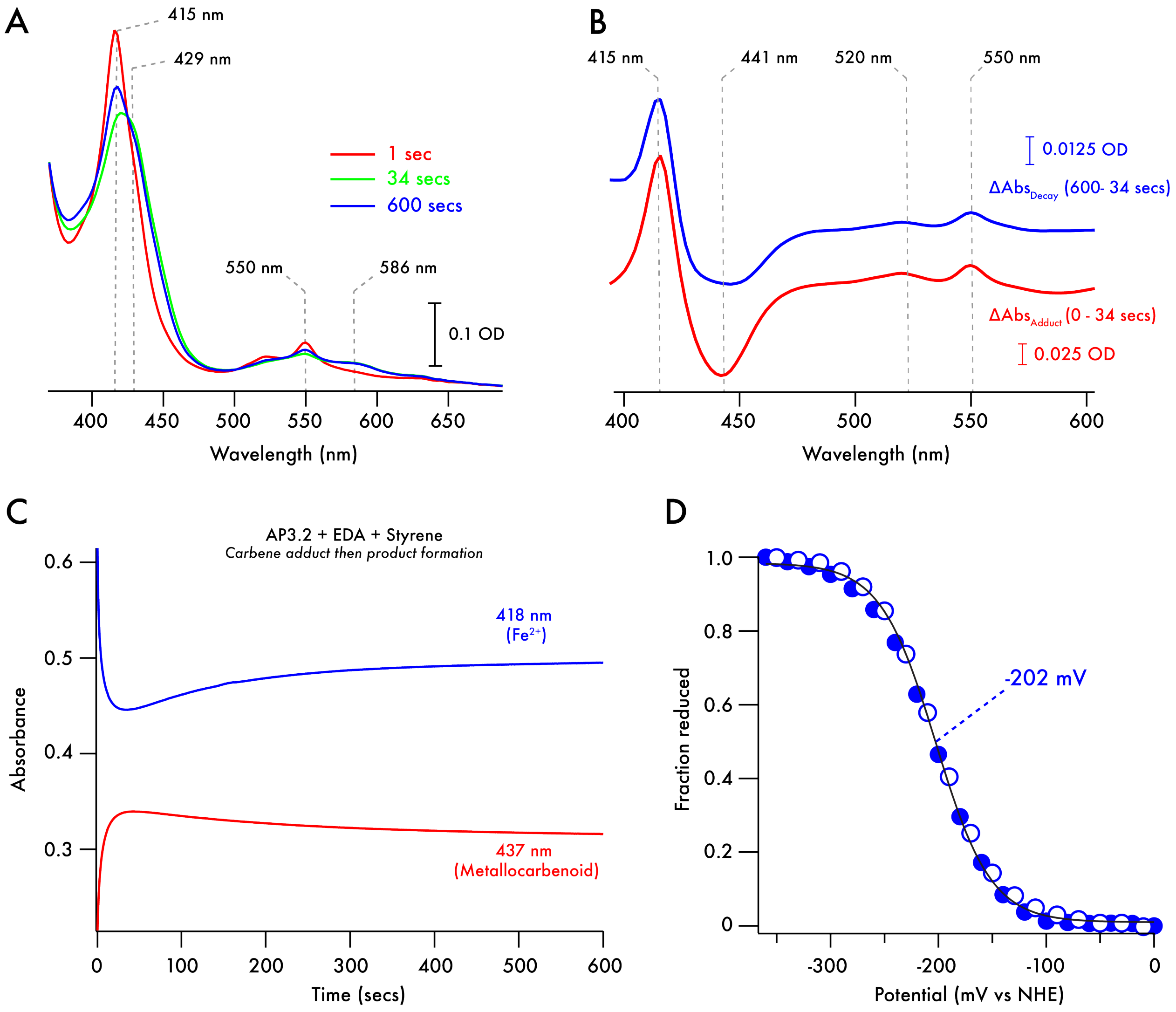


**Figure S13. Generation of the AP3.2 metallocarbenoid intermediate in the presence of 3mM styrene. A.** Electronic spectra recorded after rapid mixing of ferrous AP3.2 (7.5 µM, red trace) with EDA (500 µM) and styrene (3 mM) at 5 °C (40% EtOH) in the stopped-flow spectrophotometer. The green spectrum taken at 34 seconds post mixing decays towards the ferrous spectrum over the course of 600 seconds (blue trace). **B.** Electronic difference spectra highlighting the spectroscopic changes associated with metallocarbenoid formation and decay in AP3.2. The lower, red trace demonstrates the spectroscopic changes that occur during the formation of the metallocarbenoid, which are similar to those observed during its subsequent decay (upper, blue trace). **C.** Kinetic trace corresponding to the decay and reformation of the ferrous AP3.2 species during turnover. Data was measured at 418 nm following rapid mixing of ferrous AP3.2 with EDA in the stopped flow spectrophotometer at 5 °C. **D.** Redox potentiometry of AP3.2. Spectroelectrochemical data were collected in 100 mM KCl, 50 mM CHES, 10% glycerol, pH 8.6, with redox mediators as described in the methods. UV visible spectra were recorded during stepwise reduction and oxidation of the spectroelectrochemical cell. AP3.2 absorbance at 418 nm was converted to represent the fraction of reduced protein and plotted vs applied potential. Filled circles indicate data collected during the reductive steps, open circles indicate data collected during oxidative steps. Data were fitted to a single electron Nernst model.

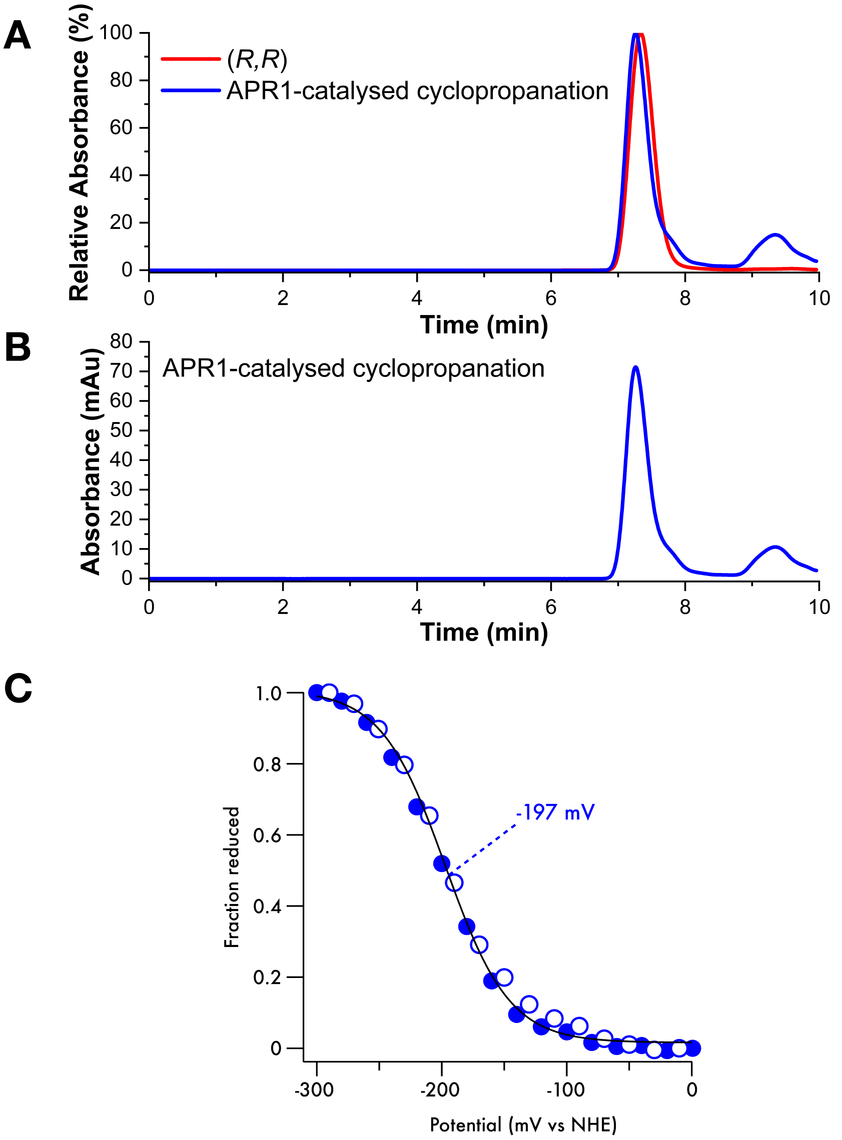


**Figure S14**. **Chiral-HPLC chromatograms for the APR1 catalyzed cyclopropanation assays. A.** Normalised APR1 (10 µM, 0.1% catalyst loading) catalyzed cyclopropanation assay between styrene (30 mM) and EDA (10 mM) (100 mM KCl, 20 mM CHES, pH 8.6, EtOH, 254 and 280 nm) (blue line) vs normalised commercial (R,R)-ethyl 2-phenylcyclopropane-1-carboxylate (red line). **B.** Average APR1 (10 µM) catalyzed cyclopropanation assay between styrene (30 mM) and EDA (10 mM) (CHES buffer, pH 8.6, 254 and 280 nm). The cyclopropane product from each assay was extracted with 1 ml of ethyl acetate and 400 µl of 3M NaOH prior to loading onto the column. A polar organic mobile phase (100% MeCN: 0.1% v/v TFA:0.1% v/v: $Et_3N$) was employed and injection volumes were 2 µl. The relative peak heights for the (R,R) and (S,S) enantiomers was used to calculate enantiomeric excess values using the equation ([R,R]-[S,S])/ ([R,R]+[S,S]). **C.** Redox potentiometry of APR1. Spectroelectrochemical data were collected in 100 mM KCl, 50 mM CHES, 10% glycerol, pH 8.6, with redox mediators as described in the methods. UV visible spectra were recorded during stepwise reduction and oxidation of the spectroelectrochemical cell. APR1 absorbance at 418 nm was converted to represent the fraction of reduced protein and plotted vs applied potential. Filled circles indicate data collected during the reductive steps, open circles indicate data collected during oxidative steps. Data were fitted to a single electron Nernst model.

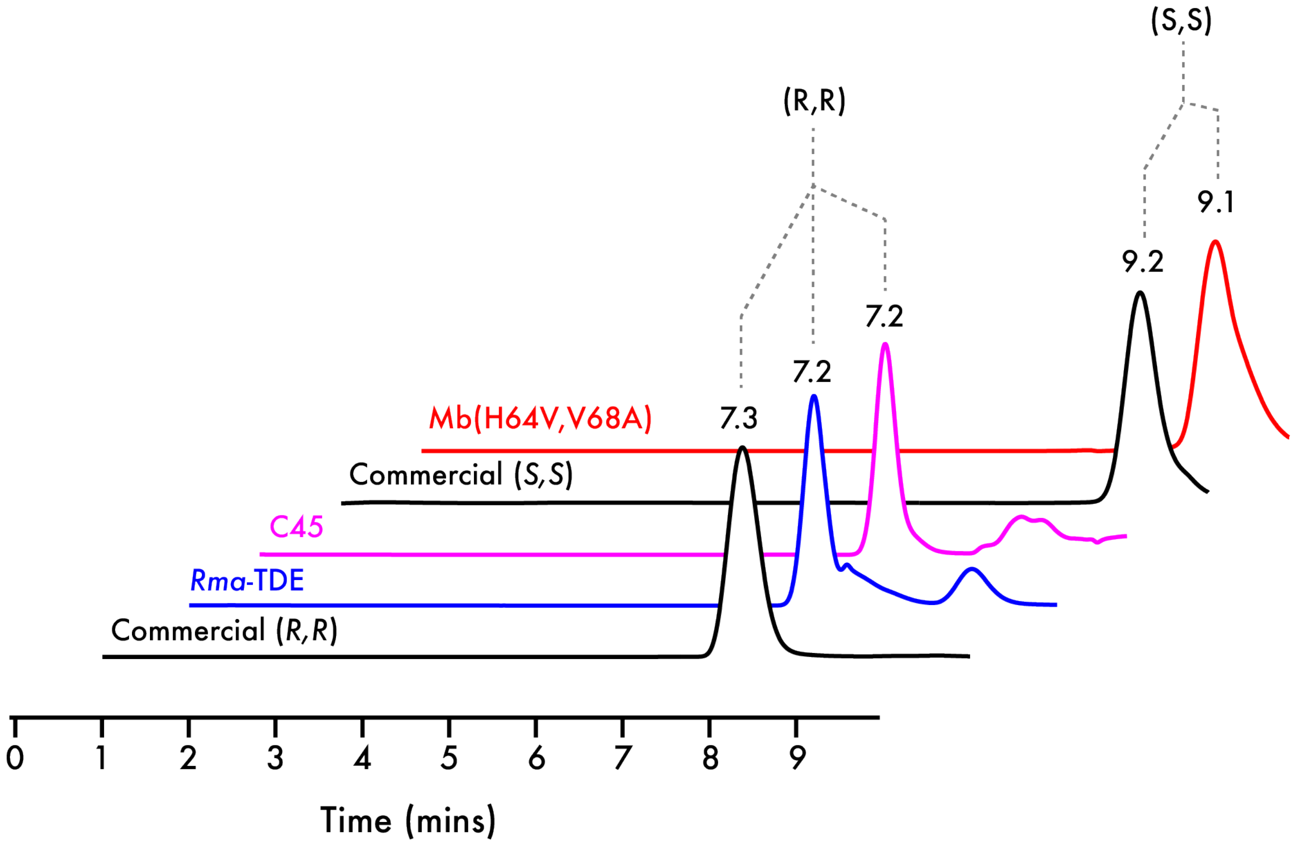


**Figure S15. Chiral-HPLC chromatograms of commercial, C45, Mb(H64V,V68A) and *Rma*-TDE catalyzed cyclopropanation assay products.** Commercial (*R,R*)-ethyl 2-phenylcyclopropane-1-carboxylate (in EtOH) and commercial (S,*S*)-ethyl 2-phenylcyclopropane-1-carboxylate (in EtOH) are represented by the labelled black traces. Also presented are chromatograms from *Rma*-TDE catalyzed cyclopropanation assay between styrene (30 mM) and EDA (10 mM) (blue trace), the C45-catalyzed cyclopropanation assay between styrene (30 mM) and EDA (10 mM) (magenta trace), and the Mb(H64V,V68A) catalyzed cyclopropanation assay between styrene (30 mM) and EDA (10 mM) (red trace). All assays were conducted in CHES buffer (100 mM KCl, 20 mM CHES, pH 8.6, 5% EtOH) and all enzyme-catalyzed reactions contained 10 µM protein (0.1% catalyst loading). The cyclopropane product from each assay was extracted with 1 ml of ethyl acetate and 400 µl of 3M NaOH prior to loading onto the column. A polar organic mobile phase (100% MeCN: 0.1% v/v TFA:0.1% v/v: $Et_3N$) was employed and injection volumes were 2 µl. All samples were recorded at 254 and 280 nm. The relative peak heights for the (R,R) and (S,S) enantiomers was used to calculate enantiomeric excess values using the equation ([R,R]-[S,S])/ ([R,R]+[S,S]).

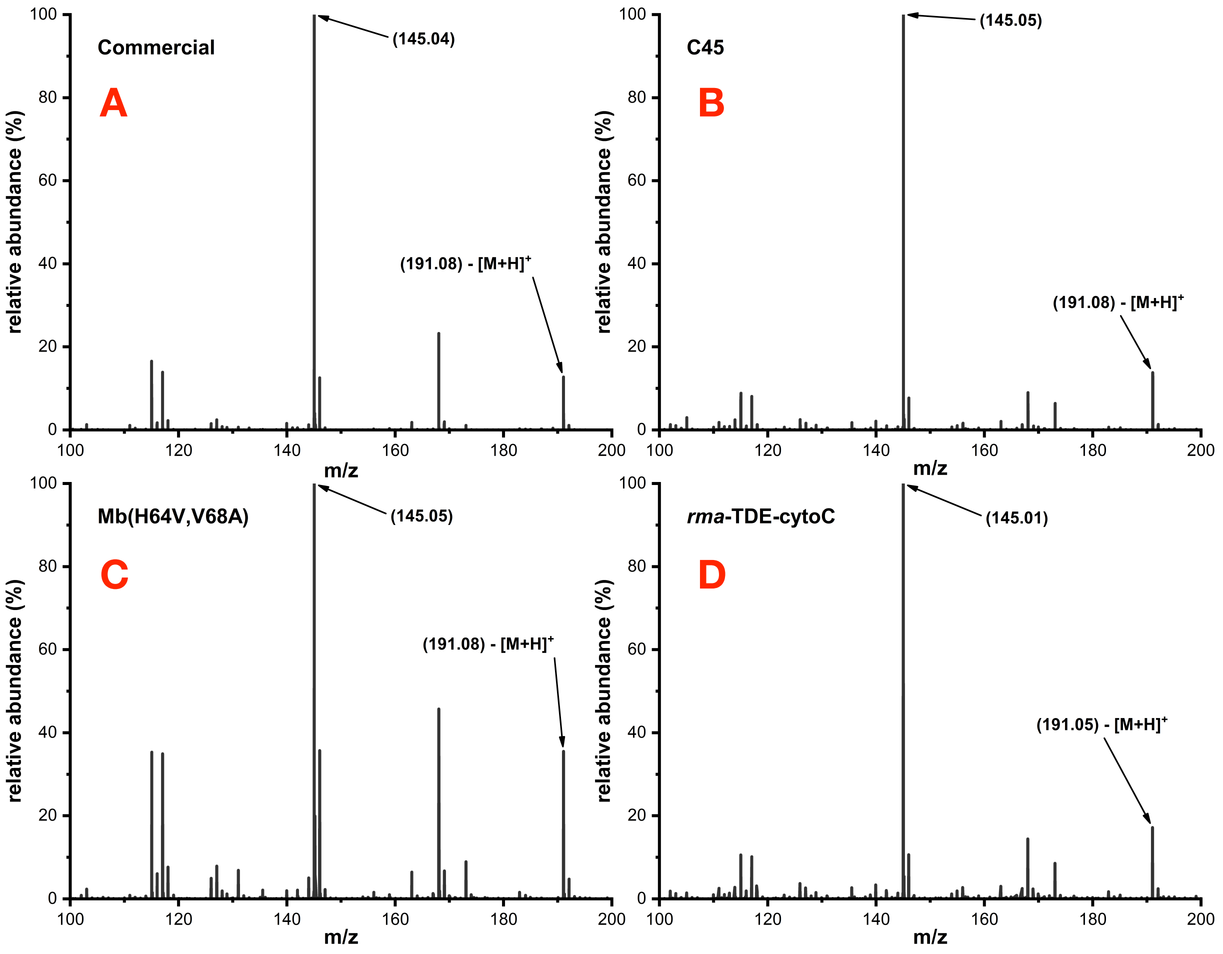


**Figure S16. LC-MS spectra of commercial, C45, Mb(H64V,V68A) and *rma*-TDE-cytoC catalyzed cyclopropanation assay products. A.** Commercial ethyl 2-phenylcyclopropane-1-carboxylate (in EtOH) exhibiting the dominant oxonium ion fragment at 145 m/z. **B.** C45-catalyzed cyclopropanation assay between styrene (30 mM) and EDA (10 mM). **C.** Mb(H64V,V68A)-catalyzed cyclopropanation assay between styrene (30 mM) and EDA (10 mM). **D.** *Rma*-TDE-catalyzed cyclopropanation assay between styrene (30 mM) and EDA (10 mM). All spectra were recorded in ES+ mode and monitored at 254 and 280 nm. A C8 column was employed for the LC separation with a gradient mobile phase (95:5:0.1% v/v water/MeCN/formate 10:90:0.1% v/v water/MeCN/formate). Assignment of major product peaks in the mass spectra (bottom).

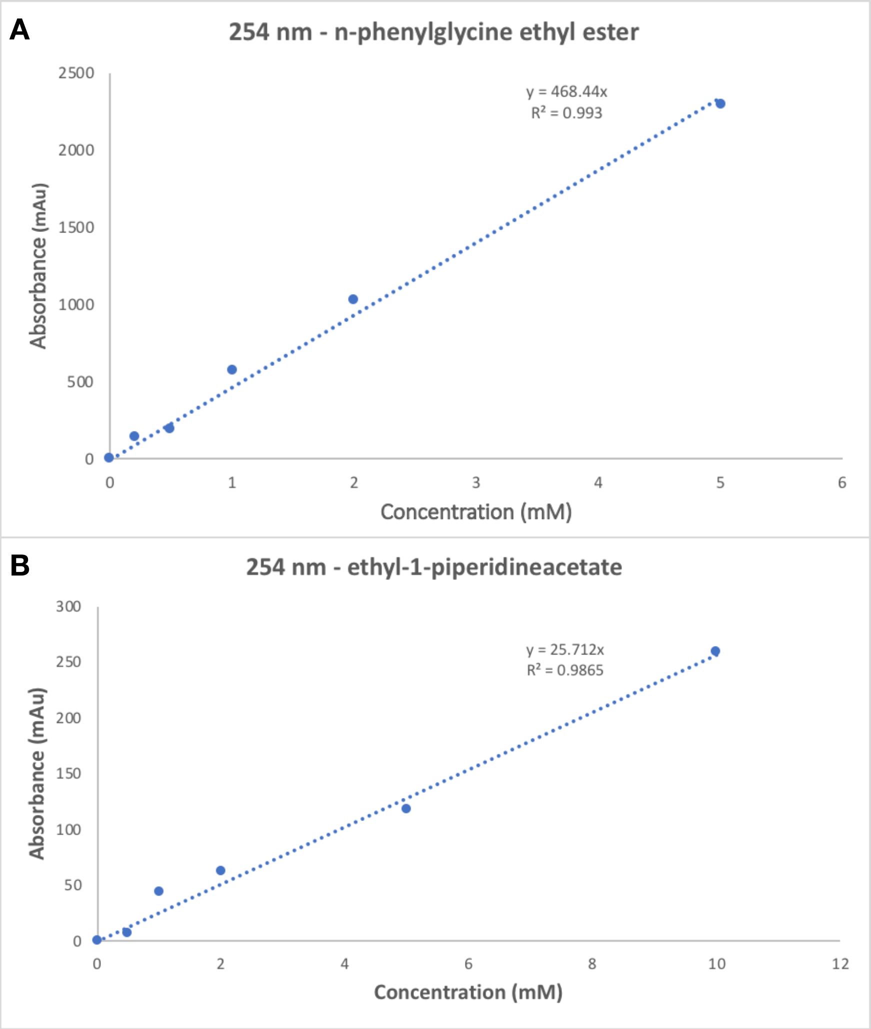


**Figure S17. Chiral-HPLC external calibrations for C45-catalyzed N-H insertion reactions. A.** Chiral-HPLC calibration for *n*-phenylglycine ethyl ester at 254 nm. A polar organic mobile phase (100% MeCN: 0.1% v/v TFA: 0.1% v/v: $Et_3N$) was employed and injection volumes were 2 µl. **B.** Chiral-HPLC calibration for ethyl-1-piperidineacetate at 254 nm. A polar organic mobile phase (100% MeCN: 0.1% v/v TFA: 0.1% v/v: $Et_3N$) was employed and injection volumes were 2 µl.

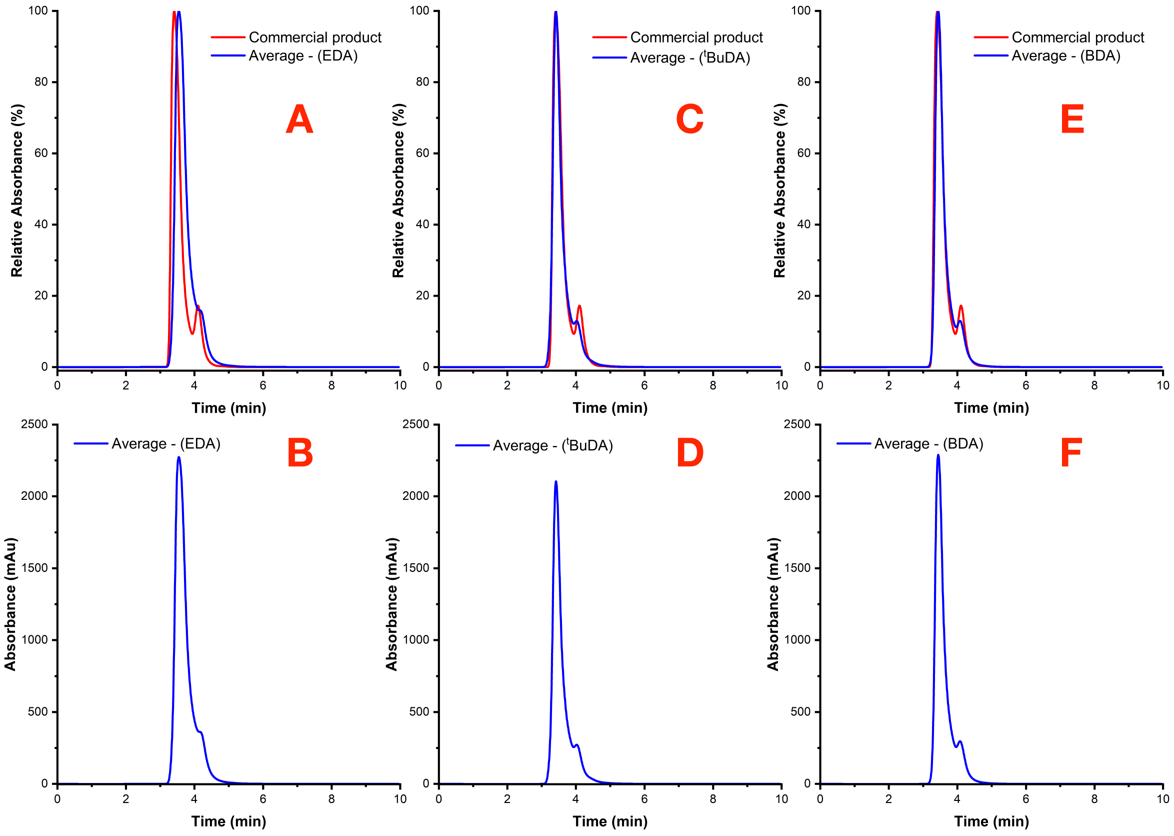


**Figure S18**. **Chiral-HPLC chromatograms for the C45-catalyzed N-H insertions reactions between *p*-chloroaniline and various diazo compounds. A.** Normalised chromatogram for a commercial sample of *n*-phenylglycine ethyl ester (red) vs normalised averaged chromatogram (blue) for the C45-catalyzed N-H insertion assay between *p*-chloroaniline (30 mM) and EDA (10 mM). **B.** Averaged chromatogram (blue) for the C45-catalyzed N-H insertion assay between *p*-chloroaniline (30 mM) and EDA (10 mM). **C.** Normalised chromatogram for a commercial sample of *n*-phenylglycine ethyl ester (red) vs normalised averaged chromatogram (blue) for the C45-catalyzed N-H insertion assay between *p*-chloroaniline (30 mM) and $^{t}$BuDA (10 mM). **D.** Averaged chromatogram (blue) for the C45-catalyzed N-H insertion assay between *p*-chloroaniline (30 mM) and $^{t}$BuDA (10 mM). **E.** Normalised chromatogram for a commercial sample of *n*-phenylglycine ethyl ester (red) vs normalised averaged chromatogram (blue) for the C45-catalyzed N-H insertion assay between *p*-chloroaniline (30 mM) and BnDA (10 mM). **F.** Averaged chromatogram (blue) for the C45-catalyzed N-H insertion assay between *p*-chloroaniline (30 mM) and BnDA (10 mM). All reactions were performed under identical conditions (100 mM KCl, 20 mM CHES, pH 8.6, 5% EtOH) with 10 µM C45 (0.1% catalyst loading). The N-H insertion product from each assay was quenched with 3 M HCl and extracted with 400 µl of ethyl acetate prior to loading onto the column. A polar organic mobile phase (100% MeCN: 0.1% v/v TFA:0.1% v/v: $Et_3N$) was employed and injection volumes were 2 µl; chromatograms were recorded at 254 nm.

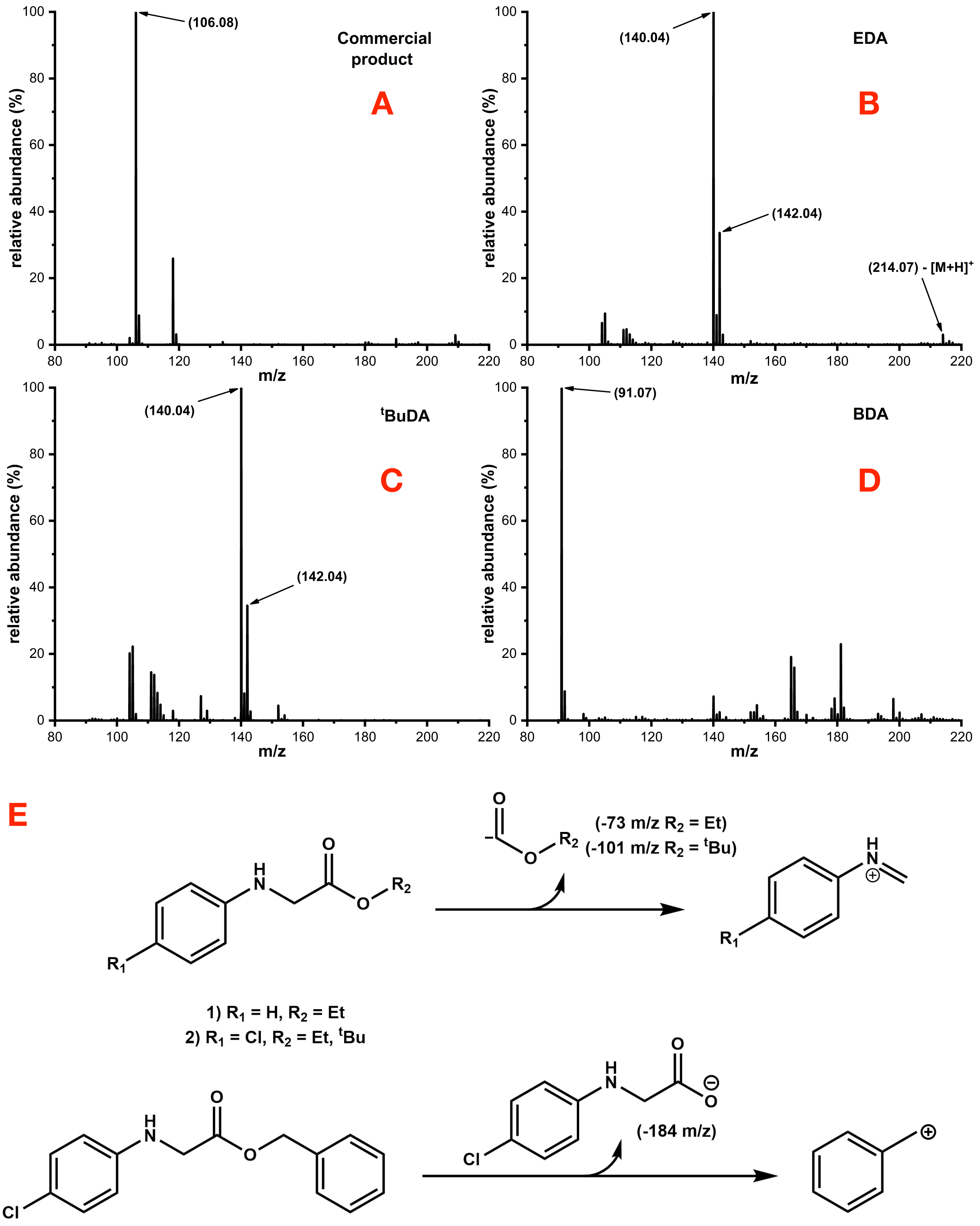


**Figure S19**. **LC-MS spectra for the C45-catalyzed N-H insertions reactions between *p*-chloroaniline and various diazo compounds. A.** A commercial sample of *n*-phenylglycine ethyl ester (in EtOH). **B.** C45-catalyzed N-H insertion assay between *p*-chloroaniline (30 mM) and EDA (10 mM). **C.** C45-catalyzed N-H insertion assay between *p*-chloroaniline (30 mM) and $^{t}$BuDA (10 mM). **D.** C45-catalyzed N-H insertion assay between *p*-chloroaniline (30 mM) and BnDA (10 mM). All spectra were recorded in ES+ mode and monitored at 254 and 280 nm. A C8 column was employed for the LC separation with a gradient mobile phase (95:5:0.1% v/v water/MeCN/formate 10:90:0.1% v/v water/MeCN/formate)**. E.** Assignment of major product peaks in the mass spectra.

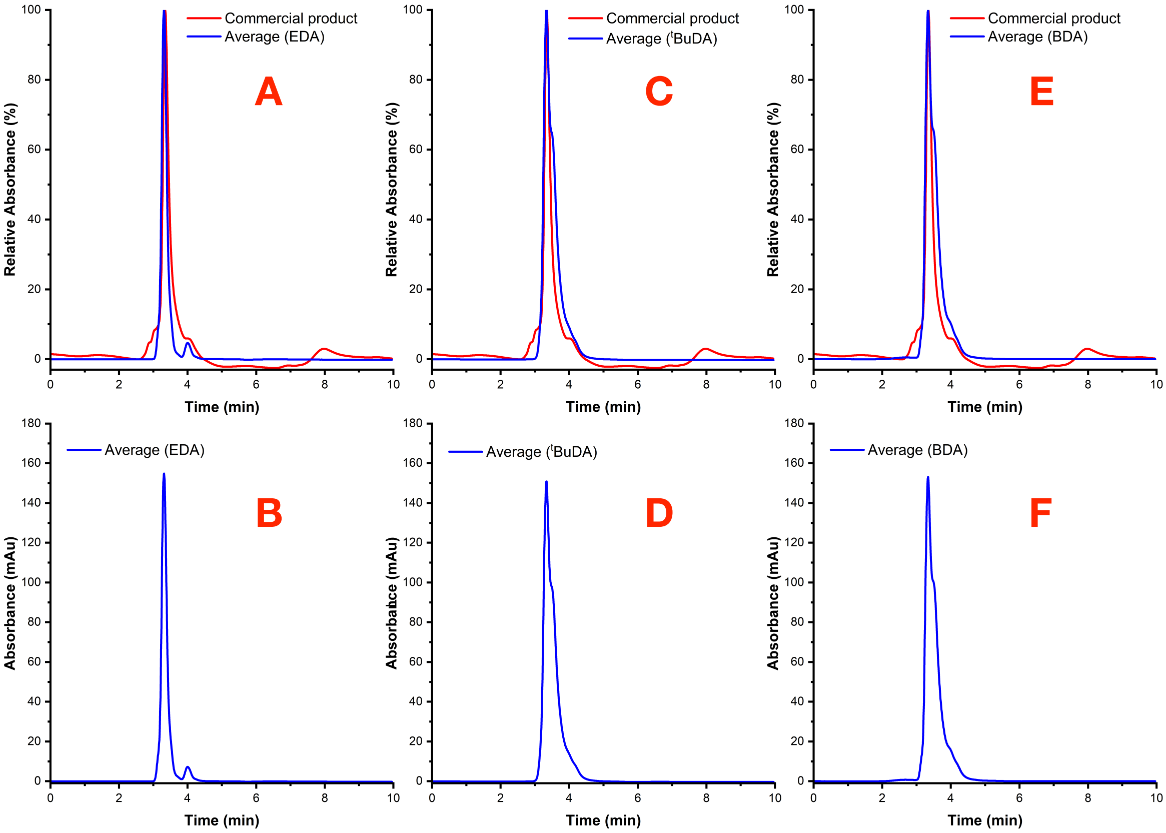


**Figure S20**. **Chiral-HPLC chromatograms for the C45-catalyzed N-H insertions reactions between piperidine and various diazo compounds. A.** Normalised chromatogram for a commercial sample of ethyl-1-piperidineacetate (red) vs normalised averaged chromatogram (blue) for the C45-catalyzed N-H insertion assay between piperidine (30 mM) and EDA (10 mM). **B.** Averaged chromatogram (blue) for the C45-catalyzed N-H insertion assay between pipieridine (30 mM) and EDA (10 mM). **C.** Normalised chromatogram for a commercial sample of ethyl-1-piperidineacetate (red) vs normalised averaged chromatogram (blue) for the C45-catalyzed N-H insertion assay between piperidine (30 mM) and $^{t}$BuDA (10 mM). **D.** Averaged chromatogram (blue) for the C45-catalyzed N-H insertion assay between piperidine (30 mM) and $^{t}$BuDA (10 mM). **E.** Normalised chromatogram for a commercial sample of ethyl-1-piperidineacetate (red) vs normalised averaged chromatogram (blue) for the C45-catalyzed N-H insertion assay between piperidine (30 mM) and BnDA (10 mM). **F.** Averaged chromatogram (blue) for the C45-catalyzed N-H insertion assay between piperidine (30 mM) and BnDA (10 mM). All reactions were performed under identical conditions (100 mM KCl, 20 mM CHES, pH 8.6, 5% EtOH) with 10 µM C45 (0.1% catalyst loading). The N-H insertion product from each assay was quenched with 3M HCl and extracted with 400 µl of ethyl acetate prior to loading onto the column. A polar organic mobile phase (100% MeCN: 0.1% v/v TFA:0.1% v/v: $Et_3N$) was employed and injection volumes were 2 µl. Chromatograms were recorded at 254 nm.

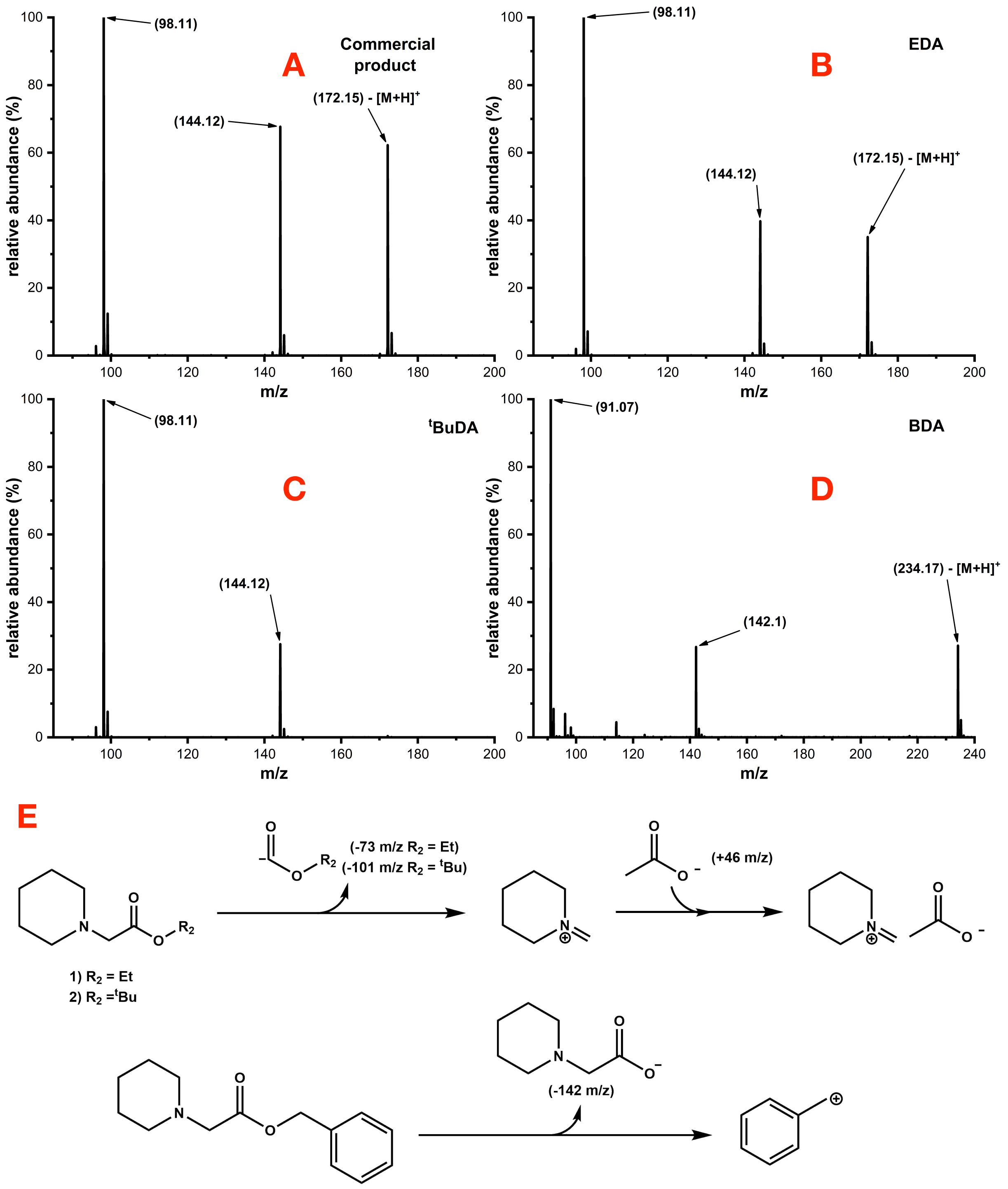


**Figure S21. LC-MS spectra for the C45-catalyzed N-H insertions reactions between p-chloroaniline and various diazo compounds. A.** A commercial sample of ethyl-1-piperidineacetate (in EtOH). **B.** C45-catalyzed N-H insertion assay between piperidine (30 mM) and EDA (10 mM). **C.** C45-catalyzed N-H insertion assay between piperidine (30 mM) and $^{t}$BuDA (10 mM). **D.** C45-catalyzed N-H insertion assay between piperidine (30 mM) and BnDA (10 mM). All spectra were recorded in ES+ mode and monitored at 254 and 280 nm. A C8 column was employed for the LC separation with a gradient mobile phase (95:5:0.1% v/v water/MeCN/formate 10:90:0.1% v/v water/MeCN/formate). **E.** Assignment of major product peaks in the mass spectra.

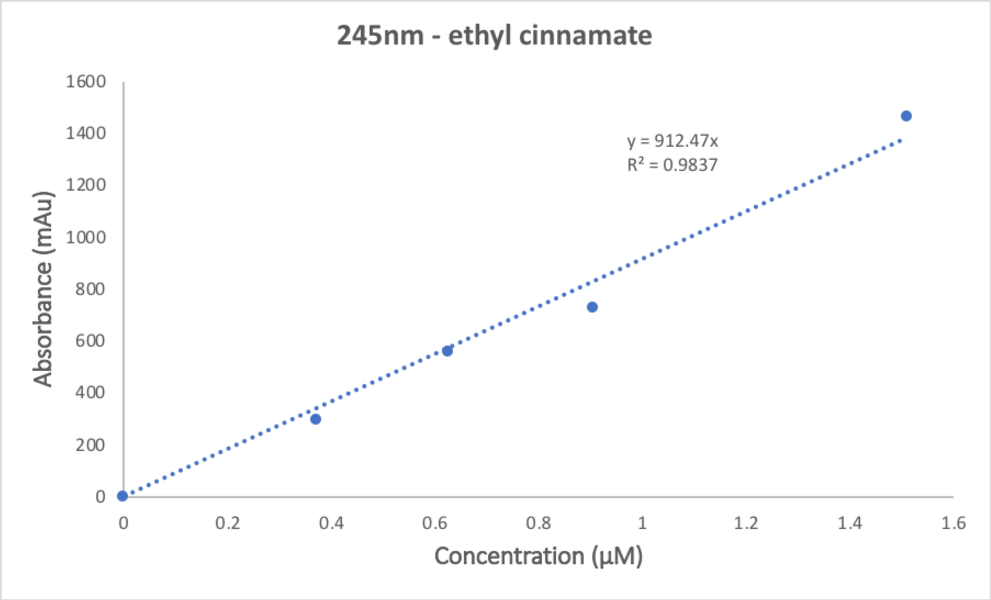


**Figure S22. C18-HPLC external calibrations for ethyl cinnamate at 245 nm.** A polar organic mobile phase (100% MeCN: 0.1% v/v TFA: 0.1% v/v: $Et_3N$) was employed and injection volumes were 20 µl.

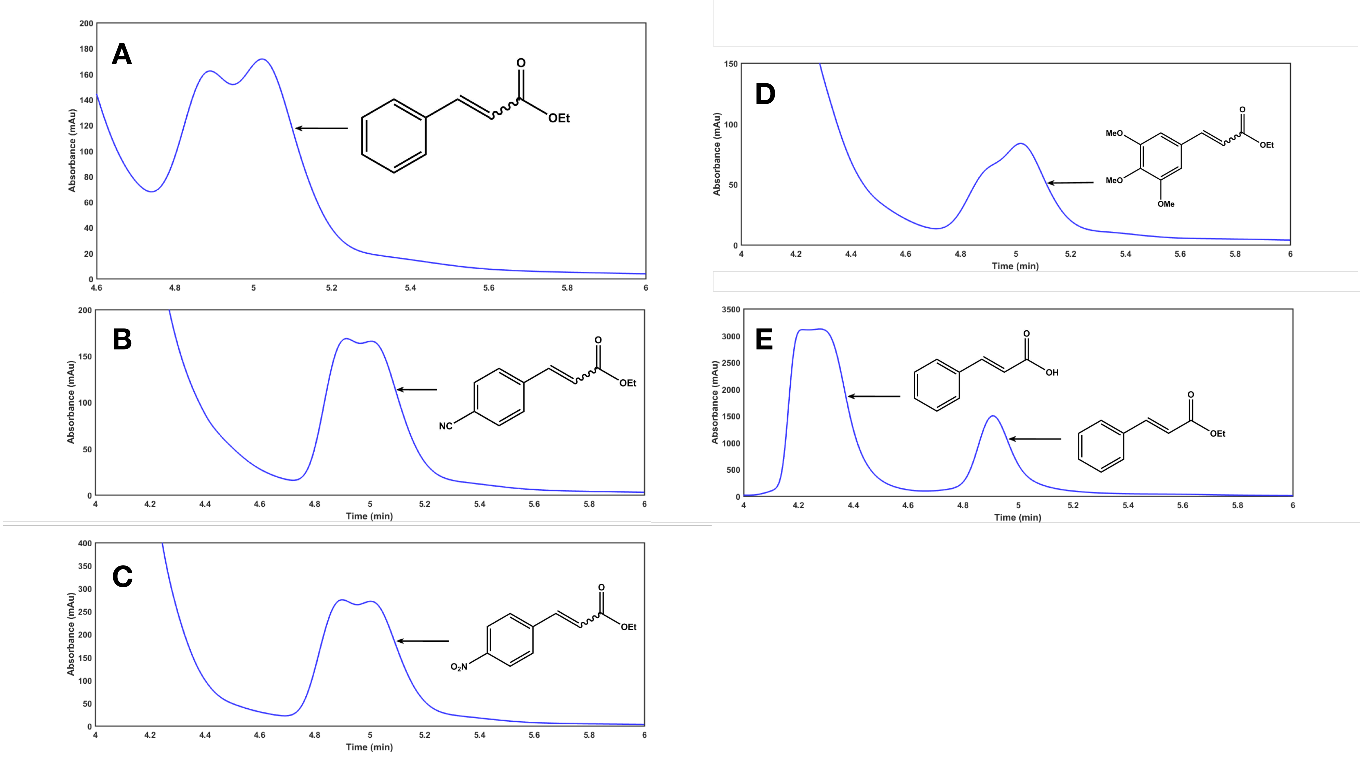


**Figure S23. C18-HPLC chromatograms for the C45 (10 µM, 0.1% catalyst loading) catalyzed carbonyl olefination assays. A.** benzaldehyde (10 mM) and EDA (10 mM), **B.** *p*-cyanobenzaldehyde (10 mM) and EDA (10 mM), **C.** *p*-nitrobenzaldehyde (10 mM) and EDA (10 mM), **D.** 3,4,5-trimethoxybenzalehyde (10 mM) and EDA (10 mM), and **E.** commercial cinnamic acid and ethyl cinnamate (in EtOH). All assays were performed in CHES buffer (pH 8.6) with 10 mM $PPh_3$ (in acetone). The α,β-unsaturated carbonyl product from each assay was extracted with 1 ml $CH_2Cl_2$ and 400 µl of 3 M NaOH prior to loading onto the column. A polar organic mobile phase (100% MeCN: 0.1% v/v TFA: 0.1% v/v: $Et_3N$) was employed and injection volumes were 20 µl; all traces were recorded at 245 nm. The *E*-isomer eluted first and was followed by the *Z*-isomer. The relative peak heights for the *E* and *Z* isomers were used to calculate the cis/trans ratio using the equation [*E*]/[*Z*].

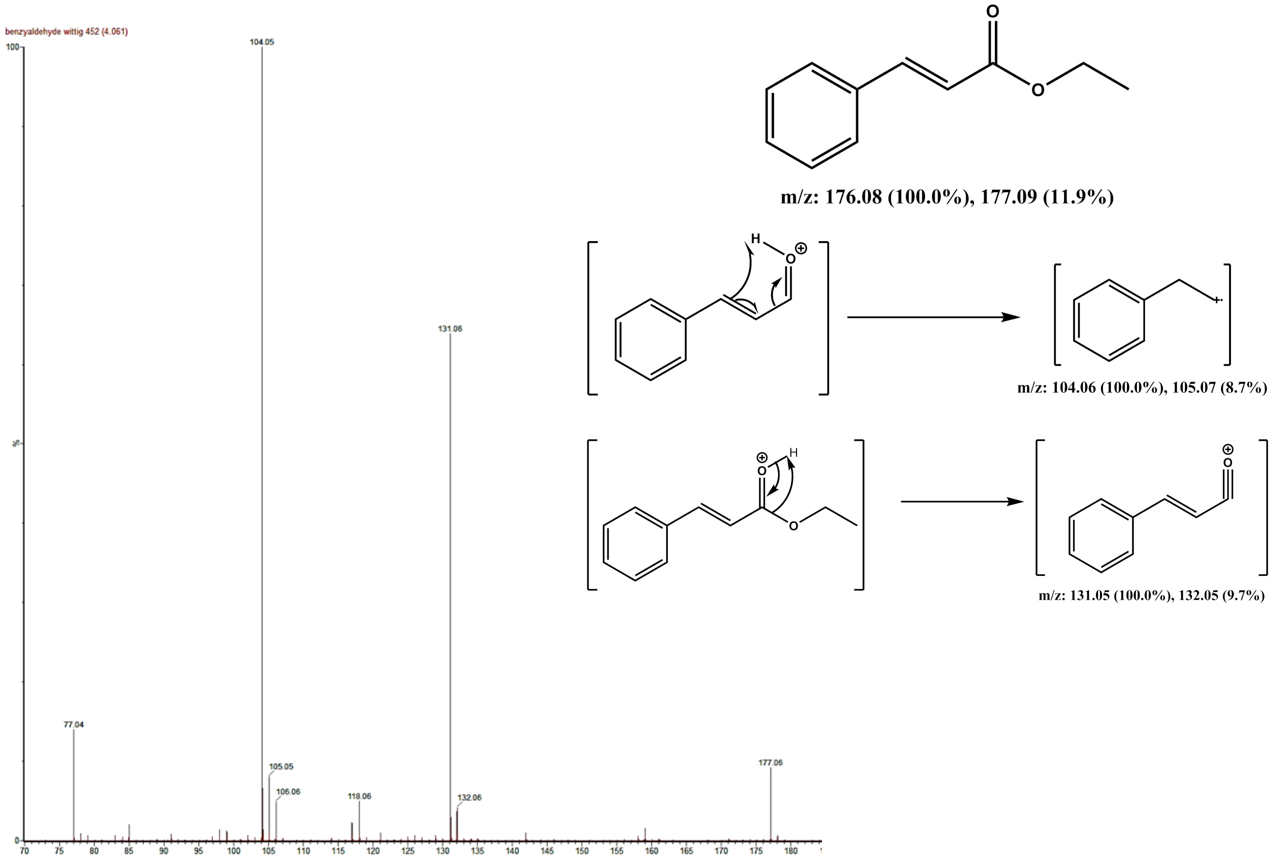


**Figure S24. LC-MS spectrum of the C45 catalyzed carbonyl olefination assay products.** LC-MS spectrum of the C45 catalyzed carbonyl olefination assay products. The mass spectrum was recorded in ES+ mode and monitored at 245 nm. A C8 column was employed for the LC separation with a gradient mobile phase (95:5% $H_2O$:MeCN to 10:90% $H_2O$:MeCN; 0.1% v/v formic acid, 0.25 ml min$^{-1}$).

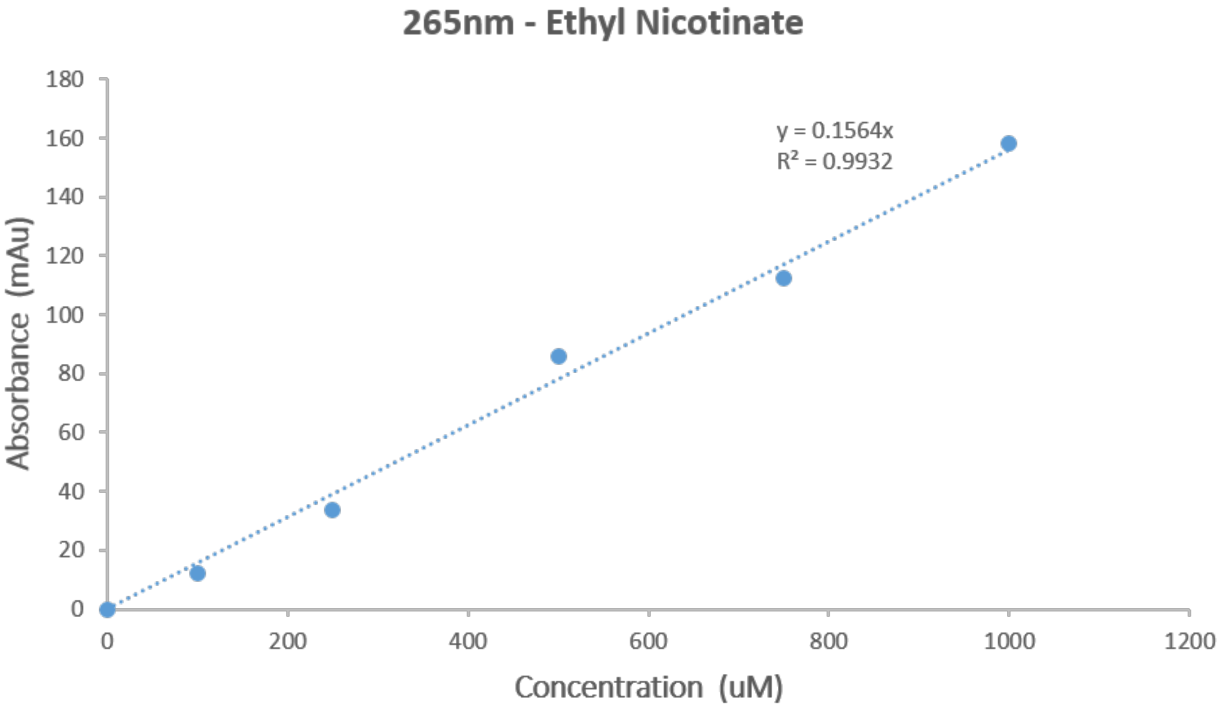


**Figure S25. Reverse phase C18 HPLC external calibrations for nicotinate at 265 nm.** A polar organic gradient was employed as the mobile phase (70:30% H2O:CH3CN to 10:90% H2O:CH3CN; 2 ml.min-1) and injection volumes were 20 µl.

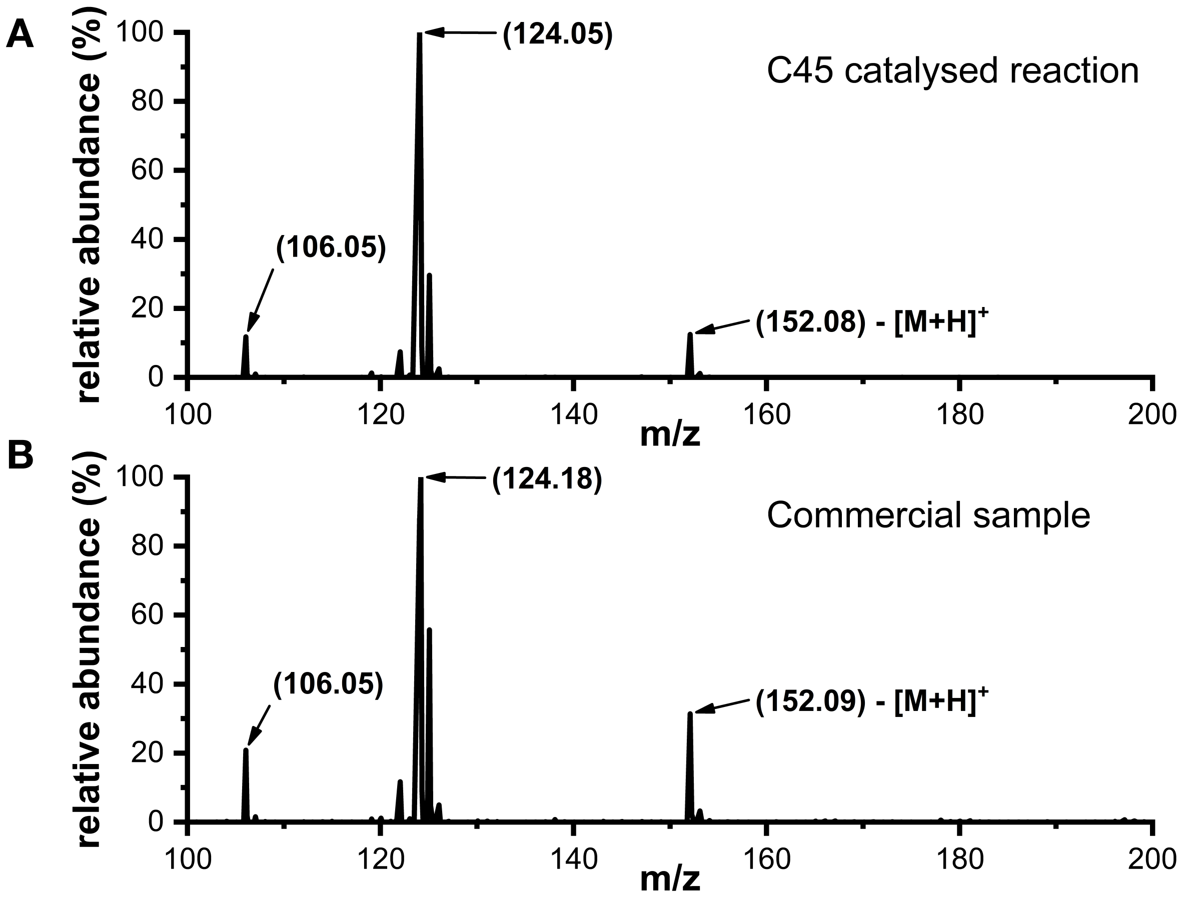


**Figure S26. LC-MS spectra of C45 catalyzed ring expansion assay product. A.** C45 (10 µM, 1% catalyst loading) catalyzed ring expansions assay between pyrrole (1 mM) and ethyl 2-bromo-2-diazoacetate (10 mM). **B.** a commercial sample of ethyl nicotinate. All spectra were recorded in ES+ mode and monitored at 265. A C8 column was employed for the LC separation with a gradient mobile phase (95:5:0.1% v/v water/MeCN/formate 10:90:0.1% v/v water/MeCN/formate)**. C.** Assignment of major product peaks in the mass spectra.

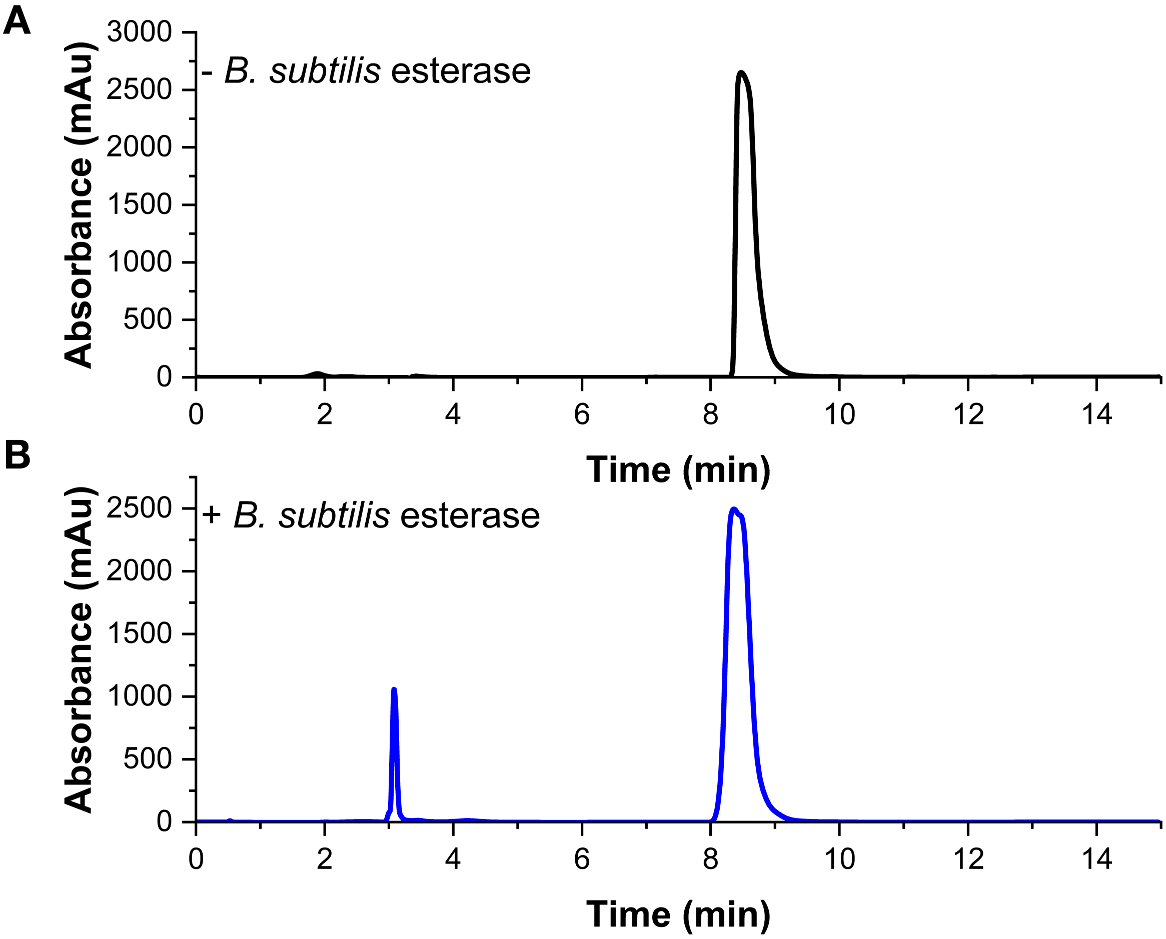


**Figure S27. C18-HPLC chromatograms of ethyl nicotinate and esterase-hydrolyzed ethyl nicotinate. A.** 50 mM commercial ethyl nicotinate (200 µl of 5 M stock in DMSO, 19.8 ml CHES buffer, pH 8.6) in the absence of esterase. **B.** 1 hour after the addition of 2 mg esterase (final esterase concentration is 100 µg/ml, 19.8ml CHES buffer, pH 8.6). The mixture was analyzed directly after precipitating the esterase with 3 M trichloroacetic acid. A reverse phase gradient mobile phase (70:30% $H_2O$:MeCN to 10:90% $H_2O$:MeCN; 2 ml.min$^{-1}$) was employed and injection volumes were 20 µl; traces were recorded at 265 nm.

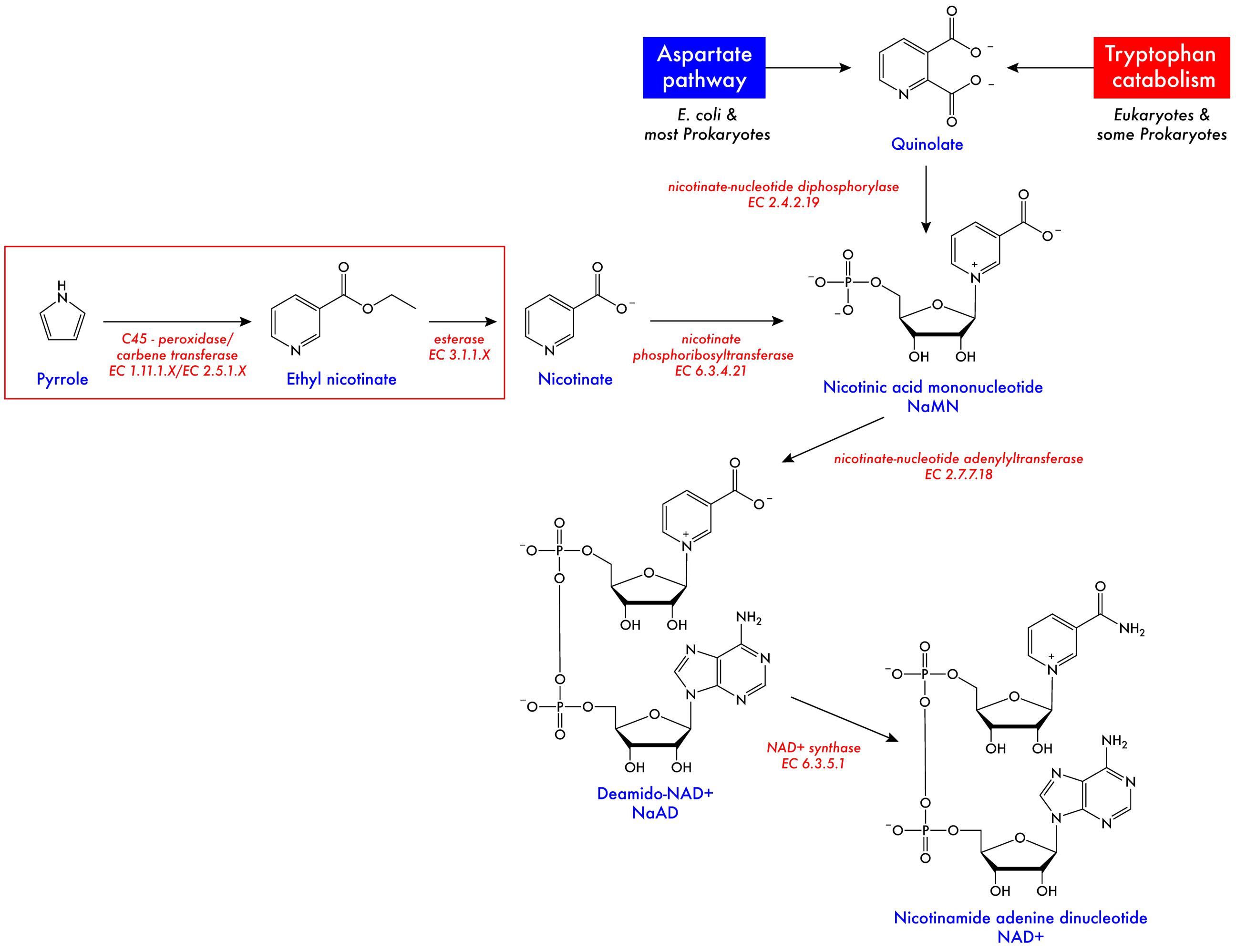


**Figure S28. Natural and engineered biosynthetic pathways to NAD+.** Steps catalyzed by the *de novo*-designed enzyme C45 and the non-native esterase from *B. subtilis* are displayed in the red box, showing an alternative route from pyrrole to nicotinate. We propose that deleting the *E. coli* nicotinate-nucleotide diphosphorylase and growing esterase- and C45-expressing cells under nicotinate starved conditions with added 2-bromo-2-diazoacetate would result in the life-sustaining biosynthesis of NAD+. Data and annotations in the figure were obtained from the KEGG database (https://www.genome.jp/kegg/kegg1.html) and the NC-IUBMB database (http://www.sbcs.qmul.ac.uk/iubmb/enzyme/).

**Table S1. C45-catalyzed cyclopropanation of styrene and *para*-substituted styrenes by EDA, and of styrene by substituted diazoacetates.** Standard deviations are quoted in parentheses.

| Product | R1 | R2 | % Yield (*R,R*) | % Yield (*S,S*) | *de* (*E*)% | *ee* (*R,R*) % | TTN (*R,R*) | Reaction time | TOF / $min^{-1}$ |
|---|---|---|---|---|---|---|---|---|---|
| **1a** | H | Et | 80.2 (±8.0) | 10.4 (±1.8) | > 99 | 77.0 (±2.8) | 802 | 120 | 6.68 |
| **1b** | OH | Et | 37.8 (±6.0) | 2.74 (±0.70) | > 99 | 86.6 (±1.3) | 378 | 120 | 3.15 |
| **1d** | $F_3C$ | Et | 91.4 (±4.7) | 0.16 (±0.04) | > 99 | 99.7 (±0.1) | 914 | 120 | 7.62 |
| **1e** | OMe | Et | 90.4 (±3.4) | 3.40 (±0.19) | > 99 | 92.8 (±0.2) | 904 | 120 | 7.53 |
| **1f** | CN | Et | 56.8 (±9.0) | 4.91 (±0.96) | > 99 | 84.2 (±0.5) | 568 | 120 | 4.73 |
| **1g** | $^t$Bu | Et | 84.6 (±1.5) | 8.22 (±0.18) | > 99 | 82.3 (±0.4) | 846 | 120 | 7.05 |
| **2a** | H | $^t$Bu | 23.0 (±10.1) | 8.72 (±1.93) | > 99 | 40.6 (±16.5) | 230 | 120 | 1.92 |
| **2b** | H | Bn | 25.1 (±1.9) | 5.15 (±1.82) | > 99 | 66.8 (±7.4) | 251 | 120 | 2.09 |

**Table S2. Comparison of enzyme-catalyzed cyclopropanations of styrene by EDA.** Standard deviations are quoted in parentheses. * indicates the Total turnover number for the specific, defined species.

| Enzyme | R1 | R2 | % Yield (*R,R*) | % Yield (*S,S*) | *de* (*E*) % | *ee* (*S,S*) % | *ee* (*R,R*) % | TTN* | Reaction time | TOF / $min^{-1}$ |
|---|---|---|---|---|---|---|---|---|---|---|
| **MgB (H64V/V68A)** | H | Et | 0 | 95.9 (±0.5) | > 99 | > 99 | - | 959 (*S,S*) | 120 | 7.99 |
| ***Rma*-TDE** | H | Et | 73.5 (±5.7) | 12.6 (±1.0) | > 99 | - | 70.6 (±2.0) | 735 (*R,R*) | 120 | 6.12 |
| **AP3.2** | H | Et | 0 | 41.5 (±4.6) | > 99 | > 99 | - | 415 (*S,S*) | 120 | 3.45 |
| **APR1** | H | Et | 85.6 (±4.3) | 11.04 (±4.8) | > 99 | - | 78.0 (±8.5) | 856 (*R,R*) | 120 | 7.13 |

**Table S3. C45-catalyzed N-H insertion reactions.** Standard deviations are quoted in parentheses.

| Product | substrate | R | % Yield (single) | % Yield (double) | single/ double | TTN | Reaction time | TOF / $min^{-1}$ |
|---|---|---|---|---|---|---|---|---|
| **4a** | piperidine | Et | 78.9 (±11.9) | n/a | n/a | 789 | 120 | 6.58 |
| **4b** | piperidine | $^{t}$Bu | 94.8 (±13.5) | n/a | n/a | 948 | 120 | 7.90 |
| **4c** | piperidine | Bn | 96.6 (±14.4) | n/a | n/a | 966 | 120 | 8.05 |
| **5a** | *p*-chloroaniline | Et | 63.1 (±3.6) | 5.3 (±5.7) | 12.1 | 631 | 120 | 5.26 |
| **5b** | *p*-chloroaniline | $^{t}$Bu | 56.0 (±0.2) | 0 | - | 460 | 120 | 4.67 |
| **5c** | *p*-chloroaniline | Bn | 60.1 (±0.2) | 0 | - | 601 | 120 | 5.01 |

**Table S4. C45-catalyzed olefination of aldehydes.**

| Product | R1 | R2 | R3 | % Yield (*cis*) | % Yield (*trans*) | *cis*/ *trans* | TTN | Reaction time | TOF / $min^{-1}$ |
|---|---|---|---|---|---|---|---|---|---|
| **3a** | H | H | H | 3.3 | 2.7 | 1.2 | 33 | 120 | 0.28 |
| **3b** | H | CN | H | 4.5 | 4.6 | 0.98 | 45 | 120 | 0.38 |
| **3c** | H | $N_2O$ | H | 7.0 | 7.0 | 1.0 | 70 | 120 | 0.58 |
| **3d** | OMe | OMe | OMe | 11.2 | 11.2 | 1.0 | 112 | 120 | 0.93 |
